# Event integration and temporal differentiation: how hierarchical knowledge emerges in hippocampal subfields through learning

**DOI:** 10.1101/2022.07.18.500527

**Authors:** Oded Bein, Lila Davachi

## Abstract

Everyday life is composed of events organized by changes in contexts, with each event containing an unfolding sequence of occurrences. A major challenge facing our memory systems is how to integrate sequential occurrences within events while also maintaining their details and avoiding over-integration across different contexts. Here we asked if and how distinct hippocampal subfields come to hierarchically, and in parallel, represent both event context and subevent occurrences with learning. Participants viewed sequential events defined as sequences of objects superimposed on shared color frames while undergoing high-resolution fMRI. Importantly, these events were repeated to induce learning. Event segmentation, as indexed by increased reaction times at event boundaries, was observed in all repetitions. Temporal memory decisions were quicker for items from the same event compared to across different events, indicating that events shaped memory behavior. With learning, hippocampal CA3 multivoxel activation patterns clustered to reflect the event context, with more clustering correlated with behavioral facilitation during event transitions. In contrast, in the dentate gyrus temporally proximal items that belonged to the same event became associated with more differentiated neural patterns. A computational model explained these results by dynamic inhibition in the dentate gyrus. Additional similarity measures support the notion that CA3 clustered representations reflect shared voxel populations, while dentate gyrus’ distinct item representations reflect different voxel populations. These findings suggest an interplay between temporal differentiation in dentate gyrus and attractor dynamics in CA3. They advance our understanding of how knowledge is structured through integration and separation across time and context.

**Significance:** A major challenge of our memory system is to integrate experiences occurring in the same context to generalize context-appropriate knowledge, while also maintaining distinct representations of these same occurrences to avoid confusion. Here, we uncover a novel mechanism for hierarchical learning in the human hippocampus that might help to resolve this tension. In the CA3 subregion of the hippocampus, the neural representations of items presented sequentially in the same context, but not in different contexts, became more overlapping with learning. In contrast, adjacent items, appearing close in time and in the same context, became increasingly more differentiated in the dentate gyrus. Thus, multiple representations in different hippocampal subregions encoded in parallel might enable simultaneous generalization and specificity in memory.

## Introduction

A challenge to a memory system is how to distinguish similar experiences that occur in the same context to avoid confusion, while also generalizing across these same experiences to extract shared context-relevant knowledge. Changes of context, termed event boundaries, are essential to constructing context-relevant knowledge because they cause us to parse continuous experience into discrete episodes, or events (Bird, 2020; Clewett et al., 2019; Clewett & Davachi, 2017; Maurer & Nadel, 2021; Radvansky & Zacks, 2017; Zacks, 2020). These events unfold in time and within each event, layered on a stable context, there is a higher frequency sequence of occurrences or subevents. For example, making breakfast entails a sequence of subevents such as brewing coffee and frying eggs, and is a distinct event from leaving our house to commute to work. As events become more familiar, such as our everyday-life routines, learning provides opportunities to integrate subevents belonging to the same event, which can improve linking information in memory (DuBrow & Davachi, 2014; Ezzyat & Davachi, 2014; Paz et al., 2010). However, integration can also blur the distinction of event details (e.g., Brown & Stern, 2013; Chanales et al., 2019; Favila et al., 2016; Wanjia et al., 2021). Thus, it remains an important and open question how the memory system might balance between context-specific integration while also maintaining detailed representations of subevents?

Prior work has pointed towards distinct computations supported by hippocampal subfields CA3 and dentate gyrus (DG) (Norman & O’Reilly, 2003; O’Reilly & McClelland, 1994; Treves & Rolls, 1994; Yassa & Stark, 2011). The hippocampus has been implicated in the representation of context, time, and sequential events (Bellmund et al., 2018, 2020; Buzsáki & Tingley, 2018; Davachi & DuBrow, 2015; Eichenbaum, 2017; O’Keefe & Nadel, 1978). In CA3, highly interconnected auto-associative networks have been hypothesized to promote attractor dynamics: similar inputs converge to the same CA3 activity pattern (‘pattern completion’), whereas different inputs are attracted towards different activity patterns (Kesner & Rolls, 2015; Knierim & Neunuebel, 2016; Lee et al., 2004; Marr, 1971; Treves & Rolls, 1994; Vazdarjanova & Guzowski, 2004; Yassa & Stark, 2011). Through this mechanism, subevents that unfold in the same context and share perceptual and temporal information can be further integrated, whereas at event boundaries, changing perceptual input may drive CA3 towards a different pattern, representing the new context (Hasselmo & Eichenbaum, 2005; Howard et al., 2005; Kesner & Rolls, 2015).

As CA3 representations of items from the same event become more similar, DG might distinguish these same sequential representations. The DG has been broadly and widely implicated in ‘pattern separation’: the allocation of distinct neural representations to highly similar information (Knierim & Neunuebel, 2016; Leutgeb & Leutgeb, 2007; Nakazawa, 2017; Treves & Rolls, 1994; Yassa & Stark, 2011). Most if not all of this work has focused on pattern separation of highly similar spatial environments (Baker et al., 2016; Berron et al., 2016; Danielson et al., 2016; Leutgeb et al., 2007; Leutgeb & Leutgeb, 2007; Neunuebel & Knierim, 2014; Wanjia et al., 2021) or object stimuli (Bakker et al., 2008; Kirwan & Stark, 2007; Lacy et al., 2011). For sequential events that evolve in time, subevents that appear *close in time* are the most similar in their temporal information and thus might require disambiguated neural representations, to minimize interference. We investigated whether the DG performs such *temporal differentiation*, and if so, is it sensitive to event structure?

Much of everyday life is composed of familiar routines. Repeated experiences allow opportunities for both integration and encoding of details, but also entail the risk of over-integration that can lead to the loss of contextually relevant knowledge. Theoretical models and some empirical work, mostly in rodents, suggest that both attractor dynamics and neural differentiation might require familiarity with spatial environments (Chanales, et al., 2017; Fernandez et al., 2023; Gill, Mizumori, & Smith, 2011; Leutgeb et al., 2004; Lever et al., 2002; Schapiro et al., 2017; Steemers et al., 2016; Wills et al., 2005). This motivated us to examine how context-based temporal differentiation and integration increase as sequential events become familiar, to promote adaptive learning.

## Materials and Methods

*Participants.* Thirty participants were included in this study (18 females, aged 18-35 years, mean age: 23.32). One additional participant was excluded due to exceeding movement (more than 3 mm within a scan, as indicated by MCFLIRT estimated mean displacement; most participants had below 1.5mm movement in all scans, one participant had one scan with 3mm displacement, our examination revealed that MCFLIRT could correct this motion to .6mm motion, which is within voxel. Hence, this participant was included). This criterion was set after motion correction was applied during preprocessing, before any functional or behavioral analysis. Two additional participants were excluded due to poor compliance with the task (*a priori* criteria of lower than 2.5SD of the group average in the temporal memory test, during the first or the second day of testing). The sample size of 30 was determined based on previous fMRI studies using similar approaches (Aly et al., 2018; Dimsdale-Zucker et al., 2018; Schlichting et al., 2015), and a behavioral study using a similar paradigm (Heusser et al., 2018). The participants were members of the New York University community, with normal or corrected-to-normal vision. They were screened to ensure they had no neurological conditions or any contraindications for MRI. The participants provided written informed consent to participate in the study and received a payment at a rate of $30 an hour for their time in the fMRI scanner, and $10 an hour for their time outside the scanner. The study was approved by the New York University Institutional Review Board.

### Materials

The stimulus set consisted of 144 gray-scale images of nameable objects on a white background, that were selected from a pool previously used in similar studies in our lab (Heusser et al., 2016, 2018). The images were resized to 350X350 pixels, such that the object occupies as much as possible of that square, and the rest of the square was white. During the list-learning phase, objects appeared in the center of a colored squared frame. The size of the color square was 600X600 pixels in total, with the object image covering 350X350 pixels in the center of the square, leaving a frame of 125 pixels around the image of the object. We used 6 colors in the study (brackets refer to the RGB values of the colors): red [255,0,0], green [51,221,0], blue [0,0,255], yellow [255,255,0], orange [255,127,0], and magenta [255,0,255], that were set based on Matlab’s 2018 default settings. The background of the screen was set to grey [128,128,128] during all tasks.

### Experimental Design

*Overview.* The experiment was divided into 2 consecutive days that were identical in their structure. On each day, participants learned three lists of objects. Each list included different objects. During list learning, each list repeated 5 times of identical and immediate repetitions. Before and after the 5 presentations of each list, all objects of that list appeared twice in random order. The random presentation after the list learning was followed by a temporal memory test for the order of the objects as they appeared during the list learning. On day 2, after the temporal memory test of the third list, participants’ memory of the object-background color association was tested, for all objects presented on both days. All phases were conducted in the scanner and were controlled by Matlab (R2018b), using Psychtoolbox3 extensions (Brainard, 1997; Kleiner et al., 2007; Pelli, 1997). On each day, participants received detailed instructions on all phases and practiced them before they entered the scanner and once more in the scanner before performing the tasks. Upon completion of all tasks on day 2, the participants left the scanner and were shortly debriefed. In the current study, we focused on fMRI data from hippocampal subfields in the list-learning phase, and behavioral data from the list learning and the temporal memory test.^1^ Below, we provide details on each of these tasks.

*List learning.* Participants learned 6 lists, 3 on each day of the experiment. In each list, participants intentionally encoded 24 gray-scale objects that appeared sequentially, embedded in a colored frame (Figure 1A). Objects were unique to each list. On each trial, participants had to visualize the object in the color of the frame and indicate by a button press whether the object-color combination is pleasing or unpleasing to them. Participants indicated by their index or middle finger of their left hand (counterbalanced across participants), using an MRI-compatible response box. The participants were informed that the decision is subjective and that there are no right or wrong answers, and that making this task will help them to remember the color in a later memory task. Each object appeared on the screen for 2 s and was followed by a 3.5 s inter-stimulus interval (ISI), and a .5 s fixation cross before the onset of the next trial. The white background and the colored frame remained on the screen continuously (i.e., during the trial and the 3.5 + .5 s ISI and fixation). Participants were asked to make their judgment when the object was still on the screen, although responses were collected also for 1 s after the object was removed from the screen, to allow late responses.

**Figure 1.**
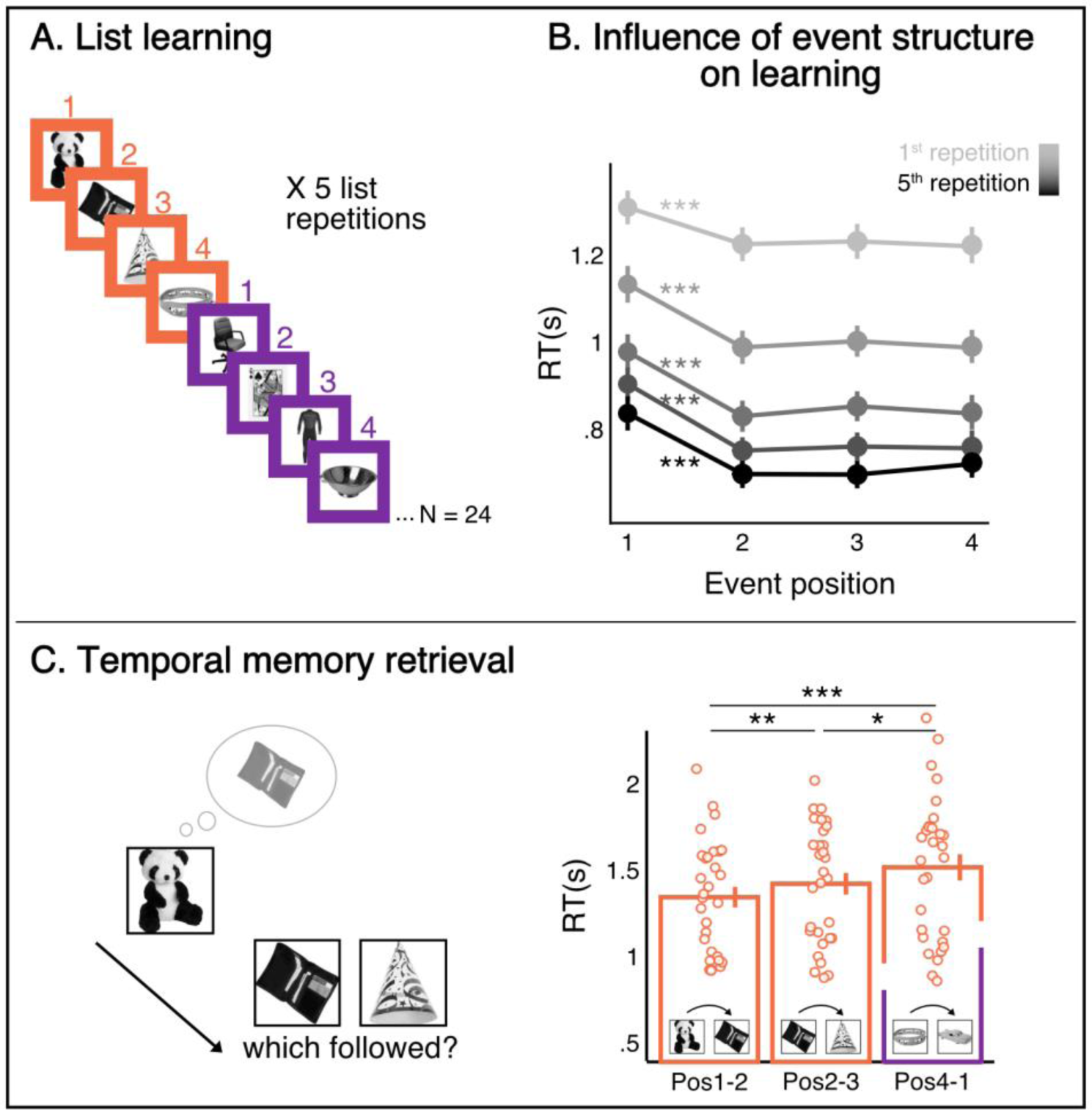
Paradigm and behavior: event structure is evident in behavior. A. During list learning, participants observed lists of 24 objects on colored frames. The color frame changed every 4 objects, creating distinct events. Each list repeated 5 times. B. Reaction times (RT) during list learning show higher RT for boundary items, a marker of event segmentation, persisting through the 5^th^ repetition. C. Left: at the end of the 5 repetitions of each list, we tested temporal order memory, by showing a cue from the list, prompting participants to retrieve the following item from the list, and then choose the correct item. Right: RT in the temporal memory test are presented for all trial types. N = 30. Data are presented as mean values, error bars reflect +/- SEM. * p = .01, ** p = .001, *** p <.0001. Statistical analysis was conducted by entering single trial RTs into general linear mixed-effects models and conducting model comparisons using Chi-square tests.

The fixed ITI during list learning was chosen for two reasons: First, different from previous fMRI studies that were designed to test specific pairs of items (e.g., DuBrow and Davachi, 2014; Ezzyat and Davachi, 2014), we designed the study look at the similarity between multiple pairs of items in a list. This was done to enable testing our main hypotheses regarding clustering of multiple items based on events, as well as the temporal differentiation analysis between different pairs of items within and across events. Altering the ITI between items would compromise our ability to compare pairs of items with different objective time gaps since it would have induced variability in both behavior and fMRI signal. We reasoned that measuring and including many pairs might be especially useful for RSA in small hippocampal subfields to overcome SNR limitations. Thus, we followed previous MVPA studies, including RSA in hippocampal subfields, that used fixed ITI (e.g., Kuhl et al., 2012; Kuhl & Chun, 2014; Kim et al., 2017), and included a rather slow ITI (4-s, along with 2-s item presentation, which makes a 6-s ISI). Second, fixed ITI allows straight-forward background connectivity analysis, because trial-evoked activity can be filtered out by using a bandpass filter that removes activation corresponding to trial frequency (see below)

The color of the frame was identical for four consecutive objects, before switching to another color for the next 4 objects. The color of the frame switched to the other color simultaneously with the presentation of the first object in an event. This made every four objects an event with a shared context, as manipulated by the frame color (Heusser et al., 2016, 2018). We refer to the first object in an event as the boundary object, appearing in event position 1, followed by non-boundary objects in event positions 2, 3, and 4. This operationalization of a sequence of items with a stable context as ‘event’ is in line with our conceptualization in the Introduction and with vast prior literature (Clewett et al., 2020; DuBrow and Davachi, 2013, 2014, 2016; Ezzyat et al., 2014; 2021; Franklin et al. 2020; Huesser et al. 2016, 2018; Sols et al., 2017). There were six events in each list, thus in total participants learned 36 events, in 6 lists. Within each list, the frame alternated between two colors (i.e., events 1, 3, and 5 were of one color, and events 2, 4, and 6 were of the other color in the colors-pair). We used pairs of highly contrasting colors to elicit clear events: red-green, yellow-blue, and orange-magenta. Each of these three color pairs appeared in one list on each day. The allocation of each color to odd-numbered events vs. even-numbered events was randomized for each participant, with the restriction that if a color appeared in odd-numbered events on the first day of the experiment, it would appear in the even-numbered events when the same color pairs appears in the second day of the experiment, to avoid having identical order of colors across lists (objects were always different across lists). The allocation of each colors-pair to the first, second, or third list on each day was also randomized per participant, with the restriction that the first list on day2 cannot include the same colors-pair as the first list on day1, to prevent participants who might have remembered the color-pairs from the first day to believe that they are going to see exactly the same experiment again on day2. To encourage associative binding of the objects, we instructed the participants to make stories linking the objects together and informed them that this would help them to correctly remember the order of the objects in the following temporal memory test (DuBrow & Davachi, 2013). The participants were also provided with an example. Critically, we did not inform or guide them regarding the different events in making the stories. Thus, for each trial, participants made judgments about the object-color combination, typically within the 2 s of object presentation (see Results), as well as created the stories, which they could do during the object presentation and the 4-s ITI (total of 6 s). In a later debrief, participants reported they have successfully performed the task and had completed their stories after a couple of repetitions. Each list was presented 5 times, immediate and identical repetition. That is, the same 24 objects (6 events) repeated in exactly the same order, with the same color frame. Each repetition was scanned in a separate scan, which began with a 2 s fixation cross (no background and color frame) and ended with a 12 s fixation cross, appearing immediately after the removal of the last object off the screen, to allow estimation of the BOLD response of the last trial.

### Temporal memory test

After the 5 repetitions of each list, participants were tested on their memory for the order of the objects (Figure 1C). On each trial, participants were first presented with an image of an object from the list, cueing them to recall the following object in the list and hold it in memory. All objects appeared without the colored frame in the memory test. The cue appeared in the center of the screen for 3 s and was followed by a 5 s white fixation cross appearing in the center of the screen. A white fixation cross was used during the delay to orient participants that they still need to hold the following object in memory. Other fixation crosses during this task were black. Then, two objects appeared on the screen next to each other, a target object and a distractor. Participants had to indicate which object immediately followed the cue object during the list learning and rate their confidence. The four response options: “left sure”, “left unsure”, “right unsure”, and “right sure” appeared below the corresponding object on the left/right, and participants indicated their response by pressing one of the corresponding 4 buttons. The participants were asked to respond as quickly as possible without sacrificing accuracy. They were further encouraged to recall the target object during the appearance of the cue and maintain it in their memory because they might not have sufficient time to recall once the two probes appear on the screen. The appearance of the target on the left or right side was randomized per participant and per list. The distractor was always the object that immediately followed the target object. We used this fine-grained test in which participants had to arbitrate between two very near objects to make sure that our lists were well-learned.

We had three types of trials: within-event boundary trial, in which the cue was the boundary object, and the target was the object in position 2 (6 trials per list, a total of 36 trials in the experiment). This is a within-event trial because both the cue and the target appeared in the same event, thus, it tests temporal memory for occurrence within a single event. As within event non-boundary trials, we had trials in which participants were cued with the object in position 2, and the target was the object that appeared in position 3 (6 trials per list, total of 36). We also had across events trials, in which participants were cued with the last object in an event (position 4), and the target item was the first object of the next event (5 trials per list, total of 30). Critically, all objects appeared with an identical temporal distance (i.e., the target immediately followed the cue) during list learning, and the objects were presented during retrieval with no color frame. Thus, the within-event and across-events trials were identical, only that in across-events trials, participants had to retrieve the target from the next event. The order of the trials was pseudo-randomized per participant and per list, such that no object repeated (as a cue, target, or distractor) with a gap of less than one trial. We also verified that the same event was not tested in any two consecutive trials (for across events trials, that meant that objects from the event that the cued belonged to, and objects from the event that the target belonged to, did not appear in the previous or next trial). The two probe objects appeared on the screen for 3 s and were followed by a 1 s fixation cross, during which responses were still collected. This was followed by an ITI of additional 3/5/7 s (average 5.25 s). Here we included jittering between trials since each trial was largely independent from other trials. To avoid variance in behavior and neural signal across different pair types, the ITI between the cue and the appearance of the targets was fixed.

during which participants performed a secondary task: an arrow appeared in the middle of the screen randomly altering between left and right every second. Participants had to indicate the direction of the arrow by a button press. This task was selected to reduce baseline activation levels in the hippocampus between trials, to improve signal estimation (Duncan et al., 2012; Stark & Squire, 2001; note that in the current study, we only report behavioral data from the temporal memory test, this optimization was done for future studies). After the arrows, there was a 1 s fixation, which, 400ms before the onset of the next trial, disappeared for 100ms, and appeared again for another 300ms. This blinking fixation was meant to prepare participants for the next trial, which began with the presentation of the cue. This task as well included a 2 s fixation cross before the beginning of the first trial, and a 13 s fixation cross after the 2 probes of the last trial appeared, to allow estimation of the BOLD response of the last trial.

### fMRI parameters and preprocessing

Participants were scanned in a 3T Siemens Magnetom Prisma scanner using a 64-channel head coil. The experiment included an MPRAGE T1-weighted anatomical scan (.9X.9X.9mm resolution) and a T2-weighted scan (.9X.9X.9mm resolution) collected at the beginning of day 1, before any task. Two pairs of fieldmap scans were acquired on each day (in each pair, one scan was acquired in the AP phase encoding direction, the other in the PA phase encoding direction). The first pair was acquired before the first list, and the second before the third list. Each repetition of each list was conducted in a single whole-brain T2*-weighted multiband EPI scan (77 volumes, TR=2000 ms, 204-mm X 204mm FOV, 136 X 136 matrix, TE=28.6, flip angle=75, phase encoding direction: anterior-posterior, GRAPPA factor 2, multiband acceleration factor of 2). In each volume, 58 slices were acquired tilted minus 20 degrees of the AC-PC, 1.5*1.5*2- mm (width*length*thickness) voxel size, no gap, in an interleaved order. At the end of day 2, participants were also given the option of running an additional MPRAGE and T2 scan. These images were collected to ensure having anatomical images in case the MPRAGE and T2 images collected on day 1 were corrupted, but unless mentioned, the day 1 images were used for the analysis.

The imaging data were preprocessed using fsl version 5.0.11 (http://www.fmrib.ox.ac.uk/fsl). Images were first corrected for B0 distortion using the acquired fieldmap scans and the topup function. For optimal correction, we corrected all the scans of each list based on the most nearing fieldmap scans. Thus, the first list on each day was corrected to the first pair of fieldmap scans, while the second and third lists on each day were corrected using the second pair of fieldmap scans. For three participants, we slightly modified the fieldmap scans we used to adapt for movement detected by the experimenter between runs, but importantly, all scans were corrected for distortion. Following distortion correction, volumes that included motion outliers were detected using fsl_motion_outliers command (framewise displacement, threshold = .9). Preprocessing was conducted using fsl’s FEAT (version 6.00). Images were highpass filtered (.1 Hz) and motion corrected within run to the middle volume using MCFLIRT. No smoothing was performed, to protect the boundaries of the hippocampal subfields, and since smoothing is unnecessary for representation similarity analysis (RSA; Kriegeskorte et al., 2006, 2008). All analyses were conducted in each participant’s native functional space and within each participant’s anatomically segmented ROIs. Note that also for the univariate analysis detailed below smoothing was redundant since we examined the average BOLD signal across all voxels in an ROI.

For the analysis of the functional data, we aligned the functional images (for RSA) and the resulting t-stats of the GLMs (for univariate analyses) in each scan to the same functional space within participant, by registering them to a template EPI image. To create this template, for each participant, we averaged together the fieldmap-corrected EPI reference images of the first scan in each day using fsl’s AnatomicalAverage command which registers and averages both images (the reference images are single volume images created by the scanner before the beginning of the scan, with identical parameters to the functional scan). This averaging across images of both days was done to avoid biasing the registration of images towards one day of the experiment. The MPRAGE image of each participant was registered to the participant’s template EPI image using fsl’s BBR registration. The transformation matrix created was then used to register the anatomical ROIs to each participant’s template EPI image, which is the participant’s native functional space.

### Regions of Interest

The hippocampal subregions, namely CA3, DG, and CA1 (in each hemisphere), as well as CSF and white-matter ROIs to be used as nuisance regressors, were automatically segmented for each participant by Freesurfer 6.0 (Iglesias et al., 2015), using both MPRAGE T1-weighted and the T2-weighted images (version 6.0 and the inclusion of the T2-weighted address the criticism raised regarding previous versions of Freesurfer, Wisse et al., 2014). We acknowledge that CA3 and DG are difficult to distinguish (Wisse et al., 2017) and that functional fMRI studies using 3T scanners have so far collapsed across these ROIs (e.g., Dimsdale-Zucker et al., 2018; Kyle, Stokes, et al., 2015; Tompary et al., 2016), preventing investigation of their different function. We leveraged freesurfer 6.0’s automatic segmentation and took a novel and automated approach to register the hippocampal subfields to each participant functional space, with the goal of retaining voxels from small hippocampal subfields to allow representational similarity analysis (RSA, see below).

The anatomically segmented ROIs are masks that include 0 or 1 values (1 marks a voxel included in that ROI). When registering these ROIs to the functional space, the alignment and resampling procedure might mix voxels together, such that voxels can have values that are between 0-1. These values reflect how much of a registered voxel originated in voxels of a value of 1, meaning, voxels that were a part of the anatomical ROI (note that registering the EPI image to the anatomical space creates the same mixture, only without knowing whether or how voxels are mixed because the original values are not given in 0 or 1. Critically, this further manipulates the functional data. Thus, we registered the anatomical ROIs to the functional space). Per each participant and each hippocampal subfield produced by Freesurfer’s anatomical segmentation, we first registered the anatomical ROI mask to the participant’s native functional space and thresholded the masks to values of .25. These registered images were then masked by a hippocampus ROI, to ensure that only hippocampus voxels were included in our subfield ROIs. The hippocampus ROI was created by combining all hippocampal subfields detected by Freesurfer, excluding the hippocampal fissure, the fimbria, and the hippocampal-amygdala transition area (HATA). Finally, for each voxel, we assigned the subfield that had the largest value, meaning that this voxel originated mostly in that subfield. We reasoned that this approach allows us to functionally dissociate hippocampal subfields (even if not fully distinguish between them) while having the obvious benefits of automatic segmentation and registration procedure, namely, being reproducible and efficient. Indeed, in both the univariate and the RSA results (see Results) we were able to dissociate CA3 from DG, demonstrating that our methodology was successful in revealing functional dissociation between these subregions.

The white-matter ROI was aligned to each functional scan, and the mean time series was extracted from each preprocessed scan using the fslmeants command. To create a confined CSF ROI (Bartoň et al., 2019), we identified, for each participant, a voxel in each hemisphere at the center of the central part of the lateral ventricle and blew a 3mm sphere around that voxel (this was done directly on the template EPI image to make sure we detect the desired voxel in the final functional space, and since the ventricles are clearly visible on EPI). We then masked that ROI with the CSF ROI created by Freesurfer, to further ensure that only CSF voxels were included. The resulting ROI was registered to each functional scan, and the mean time series was extracted from each preprocessed scan using the fslmeants command.

### Quantification and statistical analysis

Throughout the analyses of both the behavioral and the neural data, when we used mixed-level models, they were implemented using lme4 package in R (Bates et al., 2015; R core team, 2018). We used Chi-square test for statistical comparisons of the models with vs. without the effect of interest (details below for the effects included in each analysis), and AIC was used for model selection. As all our analyses are within-participant, all models included a random intercept per participant. The models further included a fixed effect of list (as factor) to capture variance related to the different lists. All analysis decisions detailed below were made *a priori* based on prior literature and recommendations. Generally, ANOVAs were *a priori* used when the measure was calculated per participant (accuracy and confidence rates, univariate fMRI analyses) whereas mixed-effect models were used when recommended to analyze single-trial data (RTs, Lo & Andrews, 2015; and RSA, Dimsdale-Zucker & Ranganath, 2018). No further statistical analyses were performed on these data.

### Behavioral data: list learning

To examine longer RT for boundaries across repetitions, we analyzed reaction time (RT) in making the pleasant/unpleasant task. No responses, too quick responses (below 100 ms), and RTs that were 3 SD above or below the average per participant, list, and repetition were excluded from the analysis. On average, 3.19% of the responses were excluded per participant. We also excluded the first response in each list, which was expectedly much higher compared to the rest of the list. Single-trial RTs were scaled without centering and entered as the predicted variable to general linear mixed models (gLMM, as implemented by the glmer function) with inverse Gaussian distribution as the linking distribution function (Lo & Andrews, 2015). We examined the effects of Boundary (boundary/non-boundary objects), Repetition (1-5 as a continuous variable), and their interaction.

### Behavioral data: temporal memory test

To test whether memory differed as a function of event position, and whether participants had to bridge across events or not in memory, we analyzed accuracy rates, high-confidence hit rates, and RT. Accuracy and high-confidence hit rates were calculated per participant and entered into a repeated-measures one-way ANOVA with Event Position (Pos1-2, Pos2-3, Pos4-1; Pos1-2, e.g., indicating that participants were cued with an object learned at event position 1, while the following object in event position 2 was the target object; Pos1-2 and Pos2-3 thus tested memory within events, while Pos4-1 trials tested memory across different events, see above). The ANOVA was implemented using the aov function in R stats package, including a within-participant error term i.e., participant/Event Position. To examine RTs, single-trial RTs of high-confidence hits were entered as the predicted variable to general linear mixed models (gLMM, as implemented by the glmer function) with inverse Gaussian distribution as the linking distribution function (Lo & Andrews, 2015). We used only high-confident hits to avoid mixing together different responses, and since high-confidence responses were the vast majority of participants’ accurate responses (see Results). The effect of Event Position was tested using all positions in one model, as well as planned comparisons examining the different pairwise comparisons between positions.

### Univariate activation

To characterize whether average BOLD signal differed between CA3 and DG during event learning, preprocessed data in each repetition in each list were entered into a voxel-based GLM as implemented by fsl’s FEAT (version 6.0). We had a regressor per each event position (4 regressors), which included all the trials in the relevant event position, modeled with 2 s boxcars locked to the onset of each trial that was convolved with FEAT’s Double-Gamma HRF function. As nuisance regressors, we included the 6 motion regressors as outputted from MCFILRT (3 for translation, 3 for rotation, in each of the x,y, and z directions of movement), the average time series for white matter, and the average time series of CSF, as well as a regressor for each time-point that was detected as outlier motion by fsl_motion_outliers (this regressor has 1 for the outlier time-point and 0 for all other time-points). To average across different lists within each participant, we registered first-level outputs to each participant’s native functional space (as detailed above) and ran a second-level (participant level) GLM that averages across lists, and thus results in a t-statistic reflecting activation per voxel per event position in each repetition.

For group-level analysis, the t-statistics were averaged per participant and per ROI and hemisphere. We quantified each hemisphere separately since previous investigations reported lateralized hippocampal effects (Bein et al., 2020; Dimsdale-Zucker et al., 2018; Kim et al., 2017; Schlichting et al., 2015). Mean activation levels were entered into a repeated measures ANOVA with the factors of Event Position (1-4), Repetition (1-5 as a continuous variable), ROI (CA3/DG), and Hemisphere (right/left), implemented by aov function in R, including a within-participant error term (participant/Event Position*Repetition*ROI*Hemisphere; note that the aov function wraps around the lm function in R). Follow-up repeated measures ANOVAs were conducted within ROI, with the factors of Position and Repetition and the relevant within participant error term. Partial eta square was estimated as effect size where relevant (sjstats package, Lüdecke, 2020).

### Item-item representational similarity analysis in time and context

To test the main hypotheses regarding learning event representations in CA3 and DG, we conducted representational similarity analysis (RSA; Kriegeskorte et al., 2006; 2008). To remove nuisance artifacts, preprocessed data was entered into a voxel-based GLM (implemented by fsl’s FEAT) with the nuisance regressors as described above: 6 motion regressors as outputted from MCFILRT (3 for translation, 3 for rotation, in each of the x,y,z directions of movement), the average timeseries for white-matter and the average time-series of CSF, as well as a regressor for each time-point that was detected as outlier motion by fsl_motion_outliers (default setting; this produces a regressor that has 1 for the outlier time point and 0 for all other time-points per each outlier time point). The residuals of this model were then registered to the template EPI, and the time course of each voxel within each ROI (CA3, DG), was extracted and z-scored (Kuhl et al., 2012; Kuhl & Chun, 2014). The third TR after the onset of each trial (starting at 4 s after the onset of the trial, capturing the peak of the HRF according to fsl’s HRF function, which is ∼4 s after the onset; Kim et al., 2017), was extracted from all voxels within an ROI to make the activity pattern corresponding to each trial. Note that this TR is before the appearance of the next trial, ensuring that patterns are not contaminated by activity related to the next trial. Additionally, due to our relatively long ISI (6 s), this time point is 10-s after the onsets of the previous trial, when most of the BOLD response for the previous trial already subsided, allowing a rather clean estimation of activation. To clean the data for RSA (as recommended in Dimsdale-Zucker & Ranganath, 2018), and particularly because correlations are susceptible to outliers, we *a priori* decided to remove voxels that demonstrated activation levels exceeding +/-3SD of the mean within each participant and each ROI (across all voxels in an ROI in all lists and repetitions) in a specific trial were *a priori* removed from all analyses involving this trial (on average across participants, 0.2% of voxel responses were excluded in each ROI).

Within ROI and participant, we then calculated Pearson’s correlation between activity patterns corresponding to each trial and the activity pattern corresponding to all other trials within a list and repetition, and Fisher transformed these correlation values. In the current study, we specifically targeted the changes in similarity due to learning, or repetition. Thus, for each correlation value, we subtracted the value of the first presentation. This allowed us to control for correlations between activity patterns that result from characteristics of the BOLD signal and processing steps (Mumford et al., 2014; Schapiro et al., 2012; Schlichting et al., 2015), which is specifically important when comparing patterns across different temporal gaps, and when the order of the trials cannot be randomized, as in our design (Mumford et al., 2014). Since the lists were identical in all repetitions, subtracting the correlation value of the first presentation from the correlation values obtained in the following repetitions alleviates the concern that our results may reflect spurious correlations. This decision was additionally informed by a preliminary step in which, to detect potential biases in similarity values, we plotted the similarity matrices across all trials in each repetition in a cerebrospinal fluid (CSF) ROI and saw some consistent biases in all repetitions. Importantly, viewing the matrices was done with all items, prior to running the main analyses comparing trials of interest. Outlier correlation values (+/-3 SD of the mean within each participant) were *a priori* excluded from the analysis following recommendations to clean data for RSA (Dimsdale-Zucker & Ranganath, 2018), and the similarity values were entered into group analysis.

The group analysis was conducted by submitting the pairwise similarity values to linear mixed-level models. We took the similarity values for temporal distances of 1/2/3 items, i.e., the similarity between an object and the immediately following object (lag of 1), as well as the next 2 objects to follow (lag of 2 and 3) in all repetitions. That is because larger distances did not occur within event, only across events. In all models, we included a fixed effect of list and a random intercept per participant.

#### Are event representations different between CA3 and DG?

In addressing this question, we treated hemispheres separately due to previous research that reported lateralized representational similarity effects in hippocampal subfields (Bein et al., 2020; Dimsdale-Zucker et al., 2018; Kim et al., 2017; Kyle, Stokes, et al., 2015; Schlichting et al., 2015; Stokes et al., 2015). Thus, the similarity values per pair of objects in repetitions 2-5 were entered into a mixed-effects linear model that examined the interaction between Event (within event/across events), ROI (CA3/DG), and Hemisphere (right/left). Since we observed a significant effect of hemisphere (see Results), we continued to test for the interaction of Event by ROI in each hemisphere. Since an interaction of Event by ROI was only found in the left hemisphere (see Results), following analyses were only conducted in the left hemisphere.

#### How do event representations change through learning within each CA3 and DG?

Within each of the left CA3 and DG ROIs, we tested the effects of Event, Temporal Distance, and Repetition, and the interaction between these factors (in the analyses that include Repetition, we included the similarity values in all repetitions 1-5 without subtracting the first presentation, as the effect of Repetition or the interaction with Repetition directly test the change across repeated presentations of the lists). To further examine how learned similarity values may change from early to later in learning, we tested the effect of Event, Temporal Distance, and their interaction in the 2^nd^ or the 5^th^ repetition. Since the observed similarity effects in both CA3 and DG were most pronounced in the 5^th^ repetition, further analyses were limited to the 5^th^ repetition. Further, in all models that did not include temporal distance as an effect of interest, we controlled for temporal distance by including this factor as an explaining variable in the model. Likewise, when testing for the effect of temporal distance, the effect of Event was included in the model to control for within vs. across events differences.

#### Are CA3 event representations related to segmentation behavior?

To foreshadow, we found an ‘event representation’ in the left CA3 in the 5^th^ repetition: higher similarity within compared to across events (see Results). We then examined whether left CA3 event representations correlated with behavioral segmentation as participants were learning the lists. To that end, we took the difference between within-event and across-events similarity in the left CA3 in the 5^th^ repetition, per participant per every two consecutive events, as well as the difference between boundary (Pos1 of the second of the two events) vs. pre-boundary RTs (the preceding Pos4, belonging to the first of the two events). These similarity differences were then entered as an explaining variable to a mixed-level model, while the RT differences were the predicted variable (linear models were used because the RT differences distribution was normal, not requiring general linear models). These difference scores control for baseline differences and examine whether similarity differences correlated with the relative increase in RTs for the boundary item. The similarity values were scaled within-participant, and as before, a fixed effect of list and event order, and a random effect per participant were added. We then followed up on this analysis by examining separately each of the values comprising these difference scores, namely, we examined whether similarity across events or similarity within event correlated with RTs of the boundary item (Pos1). Likewise, we examined whether similarity across events or similarity within event correlated with pre-boundary RTs (Pos4 of the first of each two events).

### Control for a color background effect in similarity

As mentioned above, we designed our study in a way that allows us to examine a color effect irrespective of belonging to the same sequential event, because we alternated between two background colors in each list. To examine color effects, we compared a model with a regressor noting “1” for each similarity value between items presented with the same background color and “0” for each pair of items presented with different colors, to a model without this regressor. We excluded items within the same event or across adjacent events because these might show higher or lower similarity because they belong to the same or to the adjacent sequential event, which, in turn, could artificially contribute to a color effect, given that all items in the same event share the same color, and all items from adjacent events are of different background colors. Temporal distance and list were accounted for in the model as was done above.

### Background connectivity analysis

We estimated learning-related changes in background (low frequency) connectivity between CA3 and DG. This approach was previously used to study changes in connectivity between hippocampal subfields at different phases of learning (Duncan et al., 2014; Tambini et al., 2010; Tompary et al., 2015). To remove from the BOLD signal activity directly related to items presented in the task as well as other nuisance factors, we used the residuals of the GLM that modeled univariate activation in each list and repetition of the task (see above), and which included a regressor per each event position, and nuisance regressors of csf and white-matter activity, along with 6 movement regressors (Duncan et al., 2014). These residuals, reflecting fluctuations in the BOLD signal that are not tied to items in the task, were bandpass filtered into the .01-.035 Hz range using AFNI’s 3dBandpass function (Duncan et al., 2014; Newton et al., 2011). This upper threshold is within the range of correlations between grey matter areas in fMRI (Cordes et al., 2001) and further ensures that low-frequency fluctuations do not reflect the event structure of the task, as events changed every 24 s, or .04Hz. To capture changes in connectivity due to learning, the average time course per each list in the first and last presentations (1^st^ Rep, 5^th^ Rep), was extracted per each ROI and participant, and a Pearson’s correlation was computed between the time course of CA3 and DG. The resulting correlation coefficients were Fisher-transformed and averaged across lists per each repetition. Thus, they reflect the connectivity between CA3 and DG across each repetition. We used a paired-sample t-test to compare the first presentation (1^st^ rep) and the fifth (5^th^ rep). Thus, we directly tested the difference in connectivity between repetitions, which account for baseline connectivity levels between CA3 and DG.

## Results

### Behavioral measures of learning

Participants were exposed to repeating sequences of objects superimposed on a colored frame and were asked to make judgments about the pleasantness of the object-color combination for each object. The color of the frame changed every 4 objects, representing a change in context that was operationalized as an event boundary (Figure 1A; Heusser et al., 2016, 2018). Previous studies have shown that, for once-presented events, processing time increases at event boundaries, reflecting a cost to transitioning to a new event (Heusser et al., 2018; Pettijohn & Radvansky, 2016; Zwaan, 1996). We investigated whether this marker of event segmentation persists with repetitions, indicating that participants continue to segment repeated events. Single-trial RTs for the pleasantness judgments were entered into mixed-level general linear models (Lo & Andrews, 2015), testing the effects of Boundary, Repetition, and their interaction. These models revealed a strong effect of Repetition, reflecting that participants got significantly faster over repetitions (Figure 1B), indicating learning (*χ*^2^= 3499.4, *p* < .0001, AIC reduction: 3497). We also found a main effect of Boundary (*χ*^2^= 373.87, *p* < .0001, AIC reduction: 372), indicating slower RTs for boundary compared to non-boundary items. We found no interaction of Repetition by Boundary (*χ*^2^= 0.02, *p* = .89). These results suggest that slower RTs at the boundary persisted through repetitions, as events become well learned (Figure 1B). To further demonstrate the persistence of slow RTs at the boundary across repetitions, we tested each repetition separately, and found a strong effect of Boundary in each repetition (*χ*^2^(1)‘s > 49.43, *p*’s < .0001, AIC reductions > 47). Thus, boundaries influence RTs for novel as well as familiar and well-learned events (Zwaan et al., 1995).

After learning each list, we tested temporal memory for adjacent items from the list using a serial recognition memory test. In each trial, participants were cued with an object from the list and asked to select from two options the object that came next in the list (Figure 1C). We tested temporal memory for three types of pairs: Pos1-2, Pos2-3, and Pos4-1 (the first number represents the event position of the cue, and the second number the event position of the target). As expected from presenting the sequences over 5 repetitions during learning, participants were highly accurate and confident in their responses, indicating that the lists were well learned (Accuracy: Pos1-2: *M* = 88.24, *SD* = 11.91, Pos2-3: *M* = 88.24, *SD* = 10.33, Pos4-1: *M* = 88.33, *SD* = 10.74; High-confidence hits: Pos1-2: *M* = 81.20, *SD* = 18.35, Pos2-3: *M* = 80.96, *SD* = 18.20, Pos4-1: *M* = 88.33, *SD* = 18.60, rate calculated out of total responses).^2^ No difference was observed in accuracy or high-confidence rates for the three memory trials (One-way repeated measures ANOVA with the factor of Event Position: Accuracy: *F*(2,58) = .003, *p* = .997; High-confidence: *F*(2,58) = .39, *p* = .68).

The event structure of the task was evident in participants’ RTs during retrieval (Radvansky & Zacks, 2017; Swallow et al., 2009). Interestingly, RTs were significantly slower for Pos4-1 trials, when participants had to make a temporal memory judgment for items that were from adjacent events, compared to those drawn from the same event (Pos1-2, Pos2-3; Figure 1C). Single-trial RTs were entered into a mixed-level general linear model (Lo & Andrews, 2015), which revealed a strong effect of Event Position (Pos1-2/Pos2-3/Pos4-1; *χ*^2^=30.23, *p* < .0001, AIC reduction: 26.23). Planned pairwise comparisons between specific pairs of positions revealed that RTs in Pos4-1 were slower compared to both Pos1-2 and Pos2-3 (Pos4-1 vs. Pos1-2: *χ*^2^= 31.40, *p* < .0001, AIC reduction: 29.41; Pos4-1 vs. Pos2-3: *χ*^2^ = 6.12, *p* =.013, AIC reduction: 4.12). This result suggests that the temporal order of memories that bridged an event boundary was less accessible to our participants. This is consistent with previous studies with once-presented events (DuBrow & Davachi, 2013, 2014; Ezzyat & Davachi, 2011, 2014), but extends that work to show that repeated events are still segmented in memory. We also found that RTs for Pos2-3 were longer compared to Pos1-2 (*χ*^2^ = 10.42, *p* = .0012, AIC reduction: 8.42), suggesting that cueing with boundary items benefits retrieval of items from the same event, consistent with prior work (Heusser et al., 2018; Swallow et al., 2009). We note that participants were instructed to try to recall the target item already when presented with the cue. Thus, one possibility for why we nevertheless observed RT differences is that within event, participants could recall the target already when presented with the cue (and potentially mostly when cued with the boundary), which enabled choosing the target quickly when the target and lure appeared on the screen. Across events, participants might have not been able to recall the target during cue presentation to the same extent. Thus, only when prompted with the target and lure, they had to make a memory decision, which took longer. Together, the behavioral results during list learning and in the following temporal memory test indicate that events were segmented (Hard et al., 2006; Zwaan et al., 1995).

### Univariate activation decreases with learning in DG, but not in CA3

To establish that CA3 and DG can be dissociated in human fMRI, we first examined univariate activation (average fMRI BOLD response) in DG and CA3 over the course of learning. To this end, the univariate activation estimates during learning (t-statistics resulting from a standard GLM analysis, see Methods: Univariate activation) per each event position and repetition and per each participant, averaged across all voxels in each ROI, were entered into a repeated measures ANOVA with the factors of Event Position (1 through 4), Repetition (1 through 5), ROI (CA3/DG), and Hemisphere (right/left). We found a main effect of ROI (*F*(1,29) = 7.69, *p* = .0096, η_p_ ^2^ = .21), indicating that univariate activation differed between CA3 and DG. We also found an interaction of Repetition by ROI (*F*(1,29) = 8.013, *p* = .008, η*p*^2^ = .22), demonstrating that repetition differentially modulated activity in CA3 vs. DG. A marginal effect of Event Position by Hemisphere was also observed (*F*(1,29) = 2.99, *p* = .094, η_p_ ^2^ = .09); all other effects were not significant (*F*’s(1,29) < 2.36, *p’s* > .13).

We followed up on the ROI by Repetition interaction by testing the effect of Repetition within each ROI, collapsed across hemispheres. Activation levels in each ROI were entered into a repeated measures ANOVA with Event Position and Repetition. We found a significant main effect of Repetition in DG (*F*(1,29) = 5.577, *p* = .025, η_p_ ^2^ = .16), but not in CA3 (*F*(1,29) = .34, *p* = .56). No effects of Event Position nor an interaction were observed (*F*’s(1,29) < 2.20, *p* > .14). These results reflect a decrease in univariate activation in DG from the 1^st^ repetition to the 5^th^ repetition, while CA3 activation levels remained similar, and even numerically increased (DG: Rep1: *M* = .082, *SD* = .15, Rep5: *M* = .001, *SD* = .17; CA3: Rep1: *M* = .006, *SD* = .13, Rep5: *M* = .027, *SD* = .16; average across Event Positions, the decrease in univariate activation in DG was not consistent across repetitions, see Supplementary Table 1 and Supplementary Figure 1). Thus, in our data, CA3 vs. DG revealed different activation profiles. We speculate that a lower BOLD signal in DG might reflect sparse activation, consistent with pattern separation (Leutgeb & Leutgeb, 2007; Reagh, Murray, & Yassa, 2017; Treves & Rolls, 1994; Yassa & Stark, 2011), and suggest it might not evolve linearly with learning. In CA3, activity patterns may converge towards event representation (see below), but with no changes in average activation level.

### Different event representations in the left CA3 versus DG

To test our main questions regarding neural representations in CA3 and DG, we computed the similarity between multivoxel activity patterns corresponding to sequential objects across learning repetitions (Figure 2). To specifically target changes in multivariate patterns corresponding to the learning of event structure, we used the first presentation of each list as a baseline and subtracted the similarity values for the first presentation from those measured during all subsequent presentations (Schapiro et al., 2012; Schlichting et al., 2015). This allowed us to specifically target learning-related changes in pattern similarity while controlling for correlations that might stem from the BOLD signal characteristics and analysis steps, which are identical in all repetitions (Mumford et al., 2014; Schapiro et al., 2012; Schlichting et al., 2015; Methods). Similarity values were computed for three different temporal distances (lag 1,2, and 3 items), i.e., the similarity between an object and the immediately following object (lag 1), or the object presented two or three trials later, with lag 3 being the maximum temporal distance within an event. These similarity values between objects that appeared in the same event (within event) could therefore be compared to similarity values between objects with the same temporal distance, but from different events (across events; Figure 2, Methods: Item-item representational similarity analysis in time and context). The similarity values per pair of objects were entered into a mixed-effects linear model that examined the interaction between Event (within event/across events), ROI (CA3/DG), and Hemisphere (right/left). Hemispheres were treated separately due to previously reported lateralized representational similarity effects in hippocampal subfields (Bein et al., 2020; Dimsdale-Zucker et al., 2018; Kim et al., 2017; Kyle, Stokes, et al., 2015; Schlichting et al., 2015; Stokes et al., 2015) as well as in hippocampal event representations (DuBrow & Davachi, 2013; Ezzyat & Davachi, 2014; Heusser et al., 2016; Hsieh et al., 2014). The three-way interaction between Event, ROI, and Hemisphere was significant (*χ*^2^ = 12.91, *p* = .0003, AIC reduction: 10.91), demonstrating different event representations in CA3 vs. DG that were also modulated by hemisphere. Thus, we followed up and examined the interaction of Event by ROI within each hemisphere. This analysis revealed a strong interaction of Event by ROI in the left hemisphere (*χ*^2^ = 19.27, *p* = .00001, AIC reduction: 17.27), but not in the right hemisphere (*χ*^2^ = .43, *p* = .51). Therefore, further analyses were limited to the left hemisphere. Left CA3 and DG similarity values are presented in Figures 3 and 5, where we focused on the 5^th^ repetition (see below; for all repetitions, see Supplementary Figure 2).

**Figure 2.**
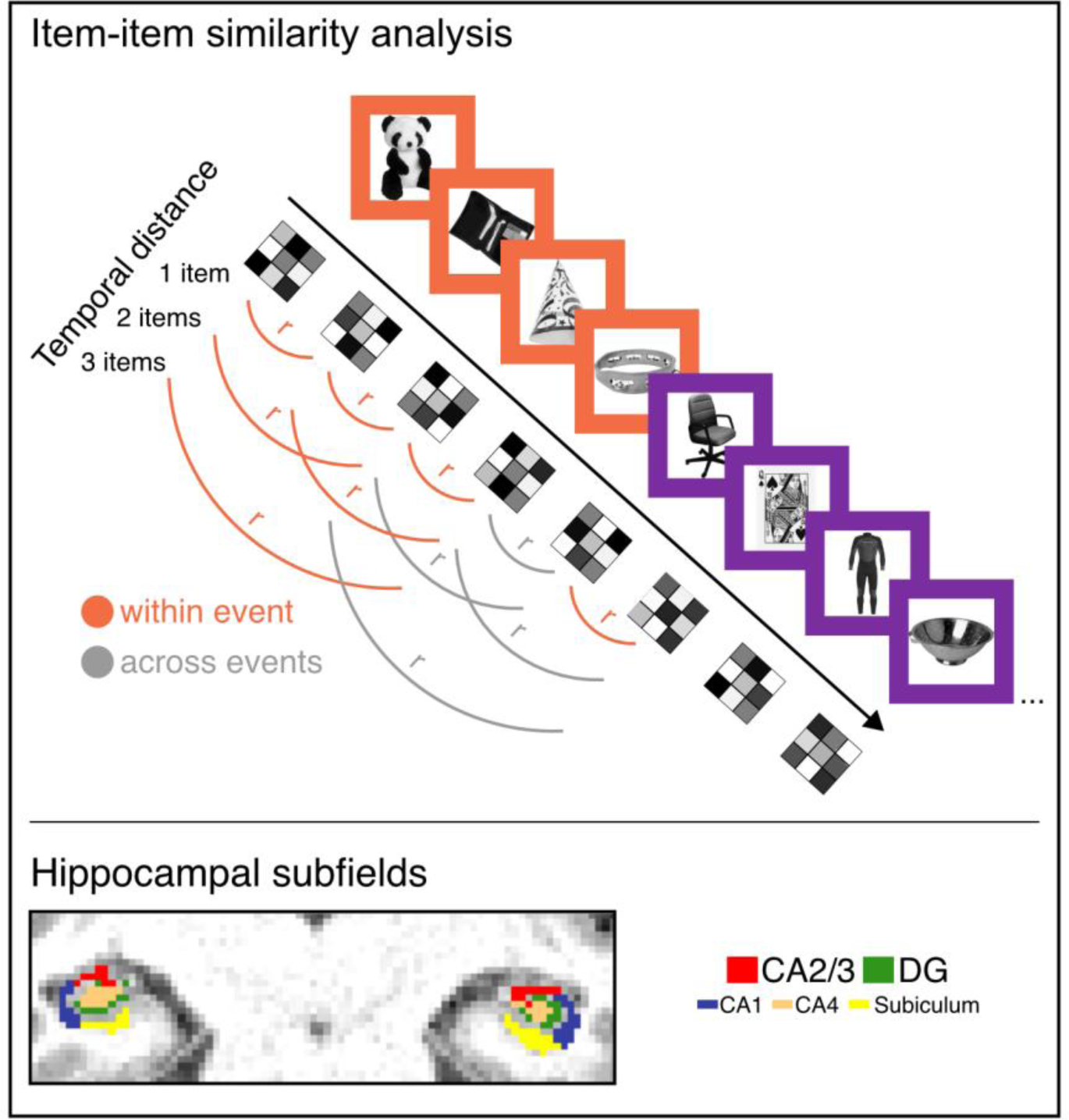
Item-item similarity in hippocampal subfields based on context and time. Top: we extracted the multivoxel activity patterns per item in the list, and computed similarity between items that belonged to the same event (within event similarity) versus items that belonged to different events (across events similarity). We also computed similarity for items that were with temporal distance of 1 item as well as temporal distance of 2 or 3 items. Bottom: patterns were extracted in hippocampal subfields, that were automatically segmented. The image shows segmentation in one participant’s anatomical space.

### Event representations in the left CA3

To further characterize the nature of the differences between CA3 and DG, we first focused on the left CA3. Following previous models of CA3 (Hasselmo & Eichenbaum, 2005; Howard et al., 2005; Kesner & Rolls, 2015), we propose that contextual stability within the same event together with a shift to a different activity pattern between events can be a mechanism by which attractor dynamics lead to integrated event representations in CA3 (Clewett et al., 2019; Davachi & DuBrow, 2015; Horner et al., 2015; Horner & Burgess, 2014; Liu et al., 2022). Our prior work has shown that the stability of hippocampal representations correlates with temporal memory for event sequences (DuBrow & Davachi, 2014; Ezzyat & Davachi, 2014). Consistent with this, it has also been shown that the magnitude of activation in CA3 during retrieval is related to holistic event retrieval success (Grande et al., 2019; Horner et al., 2015). Thus, we hypothesized that CA3 will integrate representations of information occurring within the same event while separating across different events.

Pattern similarity values for objects with all temporal distances (lag 1, 2, 3), within and across events, were entered into mixed-level models testing the effect of Event (within/across), Temporal Distance (lag 1, 2, 3), as well as their interaction (first irrespective of repetition). In CA3, we found a main effect of Event (within vs. across: *χ*^2^ = 9.06, *p* = .0026, AIC reduction: 7.06; Figure 3 and Supplementary Figure 2), with higher similarity values within, compared to across events. There was no main effect of Temporal Distance (*χ*^2^ = 2.22, *p* = .13), nor interaction effects for Event by Temporal Distance (*χ*^2^ = .04, *p* = .84). Including Repetition (1- 5) in the model yielded a significant interaction of Event by Repetition (*χ*^2^ = 4.39, *p* = .036; AIC reduction: 2.39). No significant interaction of Temporal Distance by Repetition was found (*χ*^2^(1) = .75, *p* = .39) or of Event by Temporal Distance by Repetition (*χ*^2^(1) = .0025, *p* = .96).

**Figure 3.**
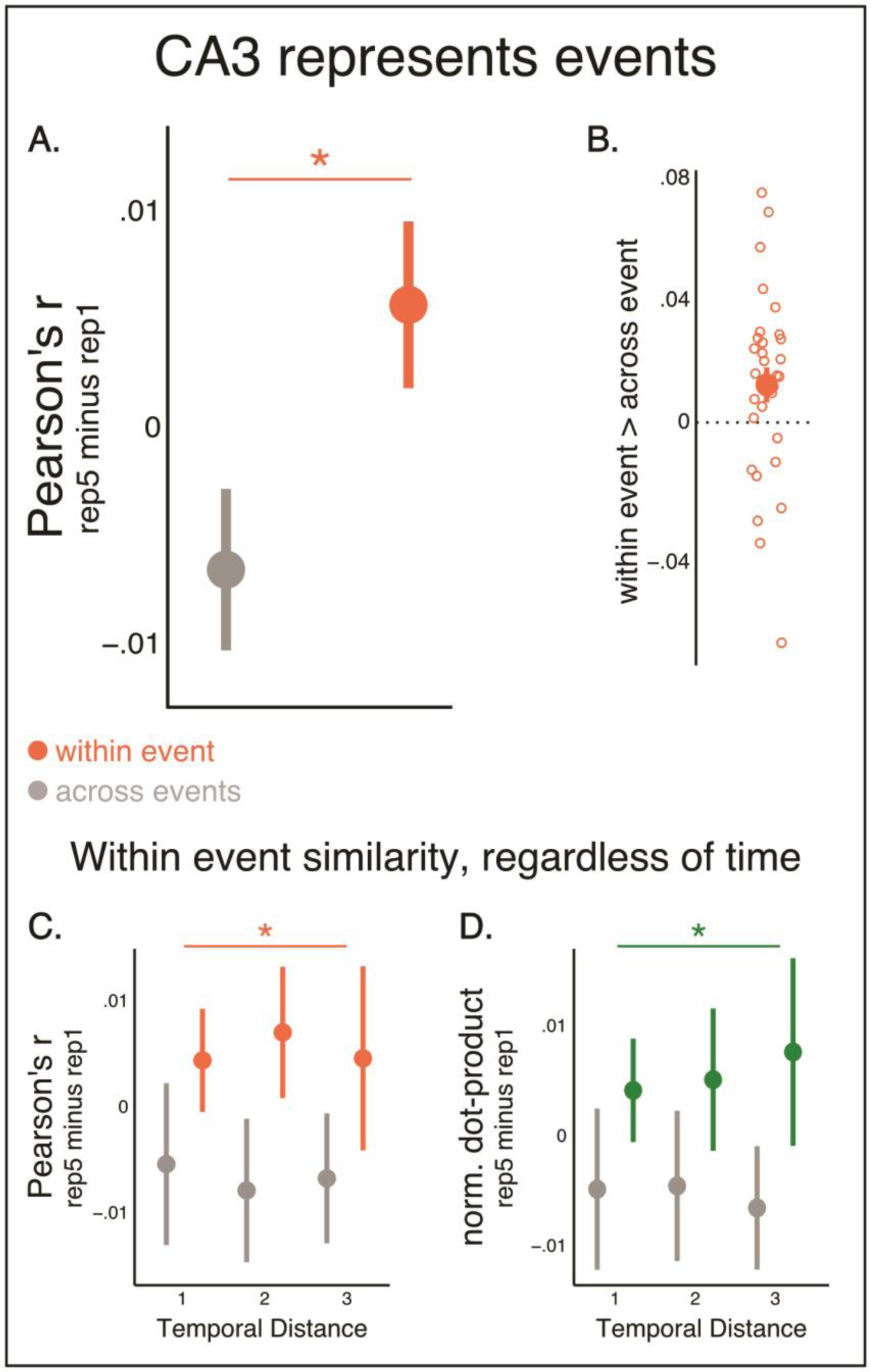
left CA3 activity patterns represent events. A. Similarity values (Pearson’s r) within event (orange) and across events (grey) in CA3. Rep5: 5^th^ repetition, Rep1: 1^st^ repetition. B. Individual participants’ values of the difference between within event and across events similarity. C. the same data as in A, but presented per each level of temporal distance. As can be seen, similarity did not differ based on temporal distance in CA3. D. Same as C but using normalized dot-product values. N = 30. Data are presented as mean values, error bars reflect +/- SEM. * p = .02, reflecting higher similarity within compared to across events, collapsed across gaps (statistical analysis was conducted by entering similarity values of individual pairs to linear mixed-effects models and conducting model comparisons using Chi-square tests, see main text).

Thus, these results revealed that CA3 representations become significantly more similar within versus across adjacent events with learning, irrespective of the temporal distance of the measured items. We next examined the 2^nd^ and 5^th^ repetitions separately, to target patterns early versus late in learning. No main effect of Event was found in the 2^nd^ repetition (*χ*^2^ = 1.55, *p* = .21). By the 5^th^ repetition, however, when events were well learned, within-event similarity was significantly higher compared to across-event similarity (*χ*^2^ = 5.42, *p* = .02, AIC reduction: 3.42, Figure 3; for additional analysis details, see Methods: How do event representations change through learning within each CA3 and DG?). Importantly, the results remained the same when accounting for univariate activation (Supplementary Information). Although not the aim of the current study, when examining similarity values for different event positions (1-4), similarity with the beginning and end of an event were numerically higher than similarity with the middle of an event (Supplementary Figure 3, see Discussion). Additionally, a control analysis confirmed that CA3 event representations are not simply explained only by items within an event sharing the same color background alone, as items from different events that shared the same background color (but were not continuous in time) were not represented similarly in CA3 (Rep5: *χ*^2^= .33, *p* = .57). Together, these results provide strong evidence for distinct event representations in hippocampal area CA3.

### CA3 event representations facilitate transitioning between events

If event representations in CA3 distinguish nearby events by allocating them to distinct neuronal ensembles, what consequences if any, might have on behavioral segmentation? As mentioned above, RTs at event boundaries items are slower compared to non-boundary items, which has previously been interpreted as a cost to transitioning between events (Heusser et al., 2018; Pettijohn & Radvansky, 2016; Zacks, 2020). We asked whether it is easier to transition to an event that is clear and distinct in its representation from the current event and thus might evoke less confusion. This could mean a relatively small cost to transitioning between these events, reflected in only a small RT increase for the boundary item. Alternatively, shifting between very different representations might render the transition to the next event more difficult, resulting in longer RTs for distinctly represented events. To examine these possibilities, we created a metric of neural clustering to index distinct representations of adjacent events by averaging similarity within each of two adjacent events and subtracting from it the similarity across these same events (see Methods: Are CA3 event representations related to segmentation behavior?). Neural clustering was then regressed with the transition cost: the difference between the RT to the boundary item in the second of these events and the immediately preceding pre-boundary item (the subtraction measure per event pairs closely controls for any baseline differences across lists or participants, see Methods). As can be seen in Figure 4, we found greater neural clustering in the left CA3 significantly correlated with less cost at the boundary, namely, a smaller RT difference between the pre-boundary and boundary items in repetition 5 (*χ*^2^ = 8.56, *p* = .003, AIC reduction: 6.56; *b* = -.40, *SD* = 13.65, reflecting that an increase of 1 SD in neural clustering led to a reduction of 40 ms, on average, in the RT difference between pre-boundary and boundary items). If neural clustering facilitates transitioning between events, it should be driven by the similarity across events, and RTs at the boundary. Indeed, lower similarity across events (more distinction) significantly correlated with faster RTs at the boundary (*χ*^2^ = 5.28, *p* = .022, AIC reduction: 3.28). Similarity within event did not explain RTs at the boundary at the pre-boundary item, nor did similarity across events explain RTs for pre-boundary items (*χ*^2^(1)’s < 1.6, *p*’s > .21). This result suggests that, for well-learned events, distinct event representations in CA3 facilitate transitioning between events.

**Figure 4.**
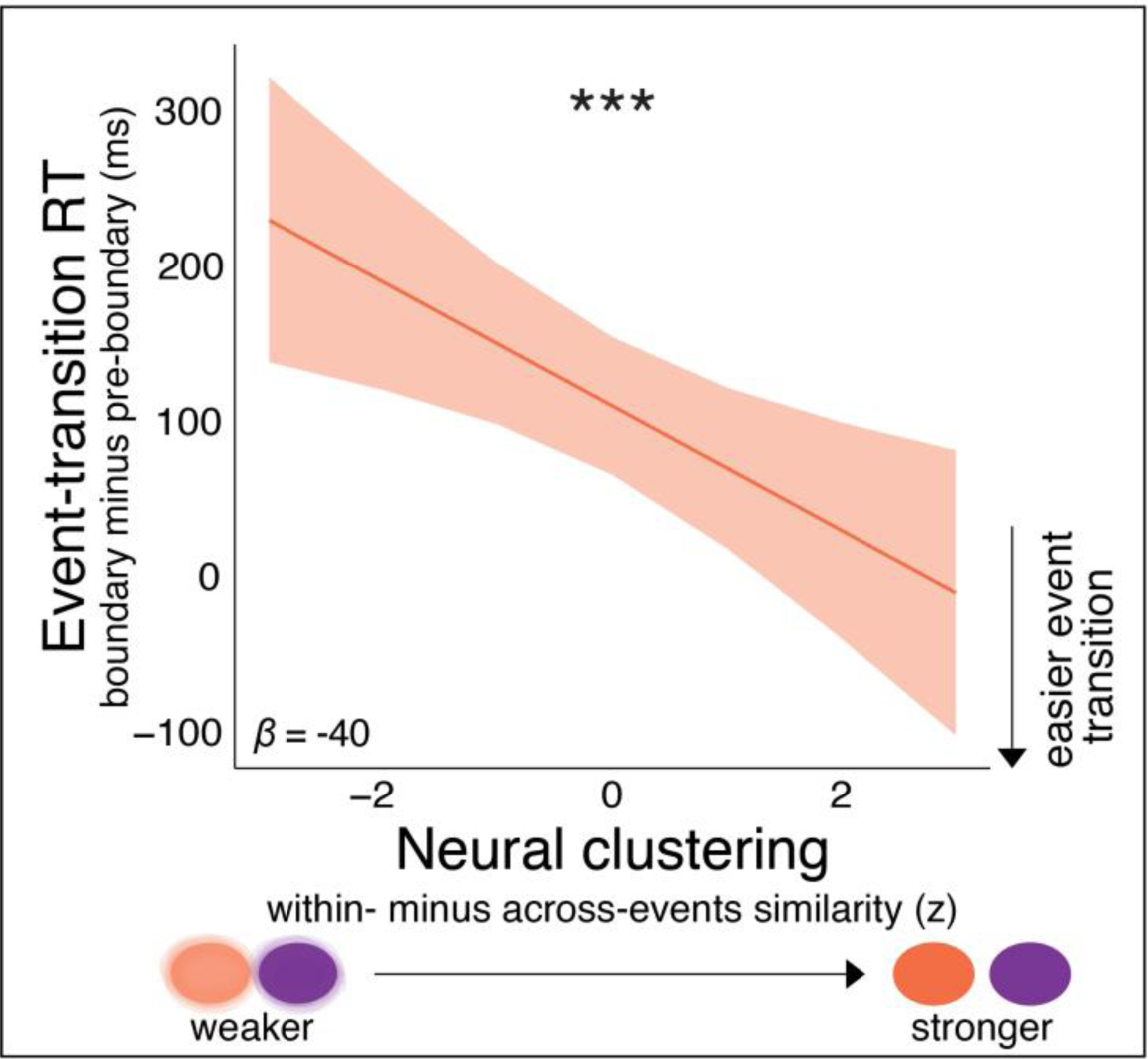
Neural clustering in the left CA3 correlates with easier event transitions. The difference between CA3 similarity within event and CA3 similarity across events (i.e., Neural clustering) correlated with an easier transitioning between events, as reflected by a smaller difference between RTs for boundary and pre-boundary items (in ms). Data are from the 5^th^ repetition. Similarity values were z-scored within participant, and the effect is plotted marginalizing over confounding factors (list, event order); thus, data are presented in SD units centered around 0. *b* = -.40, *** p = .003, results of model comparisons using Chi-square test for comparing between mixed-level models.

### CA3 representations of different events likely reflect different populations of voxels

So far, we see greater CA3 neural pattern similarity within events compared to across events and that the extent of neural event clustering facilitates event transition, as measured in response times. We next assessed the extent to which these results are consistent with attractor dynamics, in which different events are represented by different populations of CA3 neurons (Kesner & Rolls, 2015; Knierim & Neunuebel, 2016; Treves & Rolls, 1994). In fMRI, the unit of measure is a voxel so we could ask whether different voxels are engaged for distinct events. While the data reported above shows learning-related emergence of event representations, the correlation measure used (and that is prevalent in RSA studies), while capturing changes in the population of voxels activated for an item, can also be sensitive to changes in overall activation level in only a subgroup of voxels. That is because when activation levels in a subgroup of voxels changes, the mean activation level across the entire group of voxels changes correspondingly. Consequentially, the distance from the mean of each voxel changes, and with it the value of the correlation (note that this is true for Spearman’s correlation as well, not just Pearson’s correlation). Thus, we computed additional measures of representational similarity that are (1) sensitive to the overlap in the population of active voxels versus the level of activation, and (2) sensitive to changes in activation levels, regardless of the population of voxels involved.

To that end, we used two known similarity measures, normalized dot product, and the norm difference (Walther et al., 2016), that have been previously used in rodent work to examine similarity in hippocampal subfields (Madar et al., 2019). The normalized dot product is the cosine of the angle between two activity patterns (higher means more similar), normalized by their norms (the sum of squared activation values across all voxels), which makes it robust to changes in levels of activation, even in some voxels. The norm difference is the difference between the norms of two activity patterns, thus it is only sensitive to differences in the level of activation and is blind to the population of voxels contributing to that difference (see Supplementary Information). Briefly, simulations we conducted show that indeed, changing activation levels in some voxels of non-overlapping patterns influence Pearson’s correlation and the norm difference, but not the normalized dot product, while changing the overlap between patterns while holding activation levels constant influences both Pearson’s correlation and the normalized dot product, but not the norm difference (schematic description in Supplementary Figure 4, for detailed results and additional simulations, see Supplementary Information and Supplementary Figure 5).

Computing the normalized dot product and norm difference on our fMRI data, we anticipated we would replicate our correlation results in the normalized dot product measure but see no differences between the patterns’ norms. Indeed, the normalized dot product replicated our Pearson’s correlation results, finding higher similarity within compared to across events in the 5^th^ repetition (main effect of Event, within vs. across event: *χ*^2^ = 4.56, *p* = .03, AIC reduction: 2.56), with no difference based on temporal distance (main effect of Temporal Distance: *χ*^2^ = 0.017, *p* = .89; Event by Temporal Distance interaction: *χ*^2^(1) = 0.17, *p* = .68; Figure 3). No effect of Event, Temporal Distance nor an interaction was observed in the norm-difference measure (*χ*^2^’s < .60, *p*’s > .43). Note that the norm difference does not reflect average activation, but the sum of squared activation differences between activity patterns. Thus, it is unlikely that the lack of a difference stems from some voxels reducing activations, while others increasing their activation – this should have led to a major difference in norm difference. Together, the results from the normalized dot product and the norm difference measures provide further evidence that changes in univariate activation level in a subgroup of voxels are unlikely to underly low similarity between events. They are further consistent with similar populations of voxels representing items belonging to the same events, while different populations of voxels represent different events, consistent with attractor dynamics.

### Temporal differentiation in the left DG that is sensitive to context

As attractor dynamics in CA3 may pull the representations of items from the same event into a more similar space, a complementary mechanism is needed to distinguish these same sequential representations, to allow knowledge of sequential event details. For sequential events that evolve in time, subevents that appear close in time are the most similar in their temporal information and require disambiguation. In rodents, DG lesions have been shown to impair temporal memory (Morris et al., 2013), and, in humans, a prior study showed increased fMRI BOLD signal in CA3/DG in response to changes in the order of sequences of items (Azab et al., 2014), suggesting sensitivity to temporal order. However, neither of these studies has revealed how items are coded in the dentate gyrus during the unfolding of events. A recent in-vitro study found that temporally similar spike trains inputted to DG tissue resulted in divergent DG activity patterns, providing some initial evidence for pattern separation in time (Madar et al., 2019).Some recent work has provided evidence that distinct representations become revealed across learning for similar trajectories but these experiments could not dissociate time from space (Chanales, Oza, Favila, & Kuhl, 2017; Fernandez et al., 2023; Liu et al., 2022).

We investigated the following open questions: (1) Does the DG perform *temporal differentiation*? (2) if it does, is this sensitive to event structure? As above, similarity values between activity patterns of objects with temporal distances of 1, 2, and 3, both within and across events, were entered into mixed-level models. In the left DG, we found a significant interaction between Event and Temporal Distance (*χ*^2^ = 6.37, *p* = .01, AIC reduction: 4.37), stemming from lower pattern similarity between objects presented close in time, and only so if they were experienced within the same event (Figure 5, Supplementary Figure 2). Temporal Distance significantly explained variance within events, but not across events (within event: *χ*^2^ = 11.18, *p* = .0008, AIC reduction: 9.18; across events: *χ*^2^ = .14, *p* > .71). We refer to this as *temporal differentiation*, and suggest that in the left DG, temporal differentiation is specific to objects appearing within the same event, that is, the same context.

**Figure 5.**
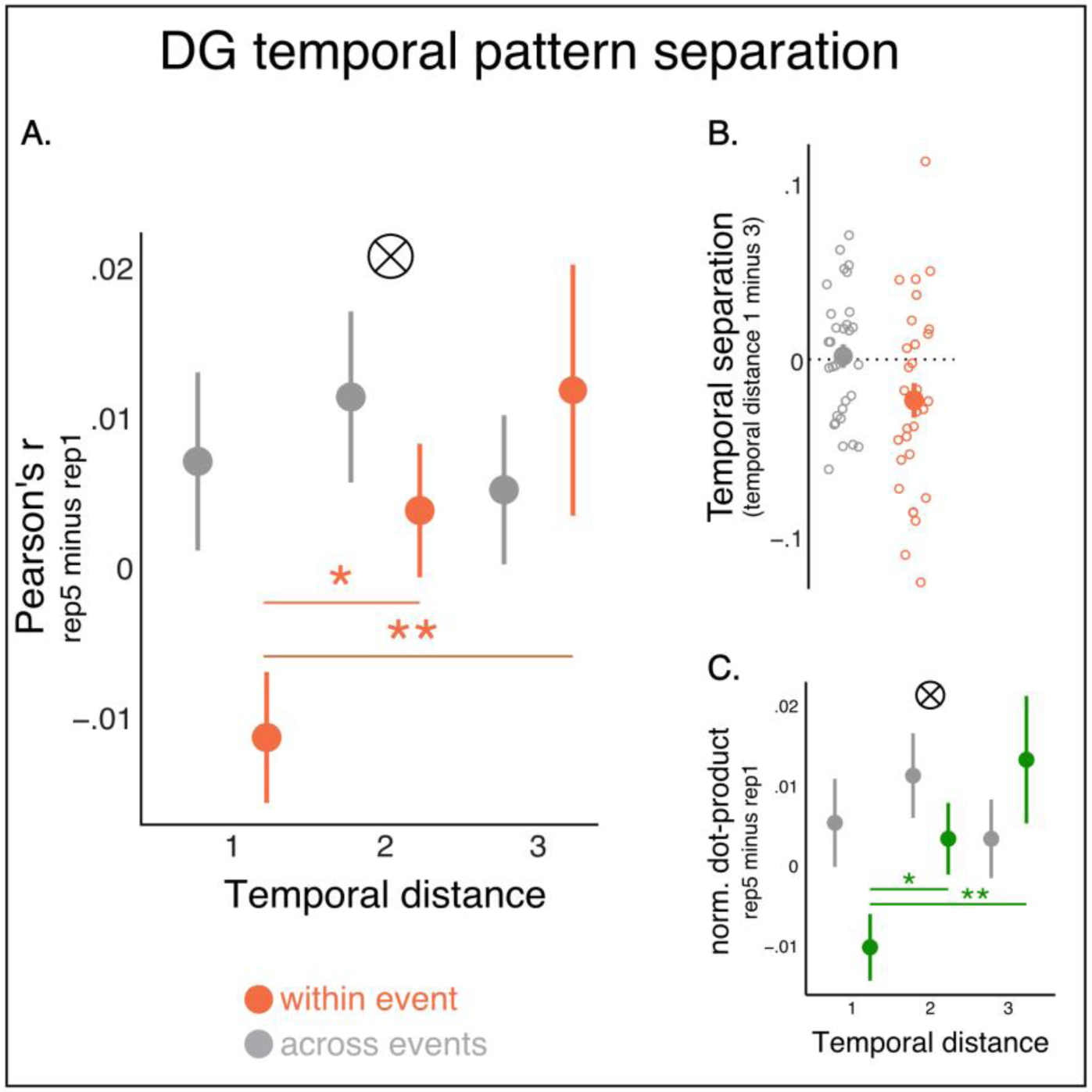
Dentate gyrus (DG) activity patterns reflect temporal pattern separation within events. A. Similarity values (Pearson correlation) in DG within event (orange) and across events (grey). Rep5: 5^th^ repetition, Rep1: 1^st^ repetition. B. Individual participants’ temporal pattern separation: the difference in pattern similarity between temporal distance of 1 object versus 3 objects, separately for within vs. across events. C. Same as A, but using the normalized dot-product values. Data are presented as mean values, error bars reflect +/- SEM. ⊗ p <= .01, Interaction of Event (within vs. across) by Temporal Distance, * p = .02, ** p = .004. Statistical analysis was conducted by entering similarity values of individual pairs to linear mixed-effects models and conducting model comparisons using Chi-square tests. N = 30.

We next examined how temporal differentiation evolved through learning. To that end, pattern similarity values from left DG were entered into a mixed-level model testing the interaction of Event, Temporal Distance, and Repetition. This revealed a significant 3-way interaction (*χ*^2^(1) = 7.17, *p* = .007, AIC reduction: 5.17), suggesting that the modulation of left DG neural pattern similarity by Temporal Distance and Event changed through learning. To better understand these changes, we followed up on this 3-way interaction by analyzing the 2^nd^ and 5^th^ repetitions separately. In the 2^nd^ repetition, we found no interaction of Event by Temporal Distance, nor a main effect of Temporal Distance, either within or across events (Interaction: *χ*^2^ = .02, *p* = .88, main effects of Temporal Distance within, across events: *χ*^2^(1)‘s < 1.1, *p*’s > .30). However, in the 5^th^ repetition, there was a significant interaction of Event by Temporal Distance (*χ*^2^ = 6.27, *p* = .01, AIC reduction: 4.27; for additional analysis details, see Methods: How do event representations change through learning within each CA3 and DG?). Pattern similarity was lower for objects close vs. far in time within event, but not across events (within event, main effect of Temporal Distance: *χ*^2^ = 10.21, *p* = .001, AIC reduction: 8.21; lag of 1 vs. 2: *χ*^2^ = 5.59, *p* = .02, AIC reduction: 3.59; lag of 1 vs. 3: *χ*^2^ = 8.11, *p* = .004, AIC reduction: 6.11; the same comparisons across events: *χ*^2^ ‘s < .23, *p*’s > .63). Further demonstrating that temporal differentiation only occurred for temporally proximal items within event, similarity was lower between items of temporal distance of 1 within event compared to across events (*χ*^2^ = 4.43, *p* = .035, AIC reduction: 2.43; the same comparison for lags of 2 and 3 was not significant: *χ*^2^ ‘s < 1, *p*’s > .30). These results remained the same accounting for univariate activation (Supplementary Information). We further note that temporal pattern separation between objects that are close in time seemed stable across the different positions in the event (Supplementary Figure 3).

Our findings show that DG differentiation is unlikely to happen between items that only share a background color, as the activity patters of items within event that shared the same background color but had a larger temporal distance (lag of 2 and 3) did not show differentiation. However, to directly examine this, we asked whether, in the 5^th^ repetition, temporal differentiation within event (computed as in the main analysis above) is stronger than differentiation between items that appeared across events with the same background color and event position (e.g., the similarity of items in event position 1 with the similarity of items in event position 2 in other events that shared the same background color, and likewise for all items in lag of 1,2, and 3). While the main analysis that compared across-events controlled for objective temporal distance, this across-event comparison reflects larger temporal distance than within events, but controls for background color and event position of items. Here as well, a significant interaction of Event (within vs. across-same color/position) and ‘Temporal Distance’ (*χ*^2^ = 4.00, *p* = .046, AIC reduction: 2.00), and no effect of Temporal Distance across-events (*χ*^2^ = 1.32, *p* = .25). This shows that temporal pattern separation (i.e., lower similarity for items that are close in time, as reported above) is specific to within-event similarity, and did not occur between items that shared the same background color but belonged to different events.

To sum up, through learning, items that were close in time and belonged to the same event, and thus potentially shared similar temporal and perceptual information, became more separated in the left DG (Kesner, 2018; Leutgeb & Leutgeb, 2007; O’Reilly & McClelland, 1994; Treves & Rolls, 1994; Yassa & Stark, 2011).

### A dynamic inhibition model for temporal pattern separation in DG

Neurophysiological literature showing uniquely high levels of inhibition in DG (Chawla et al., 2005; Coulter & Carlson, 2007; Freund & Buzsáki, 1998; Jinde et al., 2013) that determine the temporal dynamics of neuronal firing (Bartos et al., 2007). Thus, we suggest that inhibitory dynamics may account for the lower similarity between temporally adjacent items (Hasselmo & Wyble, 1997; Kesner & Rolls, 2015; Myers & Scharfman, 2011). Specifically, we propose the DG neurons that are activated for one item (e.g., item n) become inhibited for some time such that they are not active when the immediately following item (n+1) appears. This results in other neurons representing the n+1 item, and in a low overlap between activity patterns of temporally adjacent items (items n and n+1). Since inhibition gradually decays in time, for the following item (n+2), some of the neurons can overcome the lower level of inhibition and are active again, resulting in a slightly higher correlation between items n and n+2. By item n+3, inhibition decays enough such that similarity levels are high again and reach the levels of between-event similarity. Between events, this inhibition between DG neurons does not come into play because items can be distinguished based on perceptual features alone, thus DG pattern separation is unnecessary (see Discussion).

We formalized this idea using simulations, increasing the level of maximal inhibition (between items n and n+1) across repetitions, but keeping the decay in time parameter constant. Across a range of ROI sizes and rates of inhibition decay, our simulations showed that indeed decaying inhibition results in the pattern of results observed in DG, namely, low similarity for temporally proximal items, with similarity increasing as temporal distance increases (see Supplementary Information, Supplementary Figure 6 for a schematic of the model, and Supplementary Figures 7 and 8 for full results). Conceptually, learning in such a model can be seen as shaping an activity pattern for an item that is differentiated enough from previous items, such that it defeats increasing levels of inhibition.

### DG temporal differentiation likely reflects different populations of voxels

The dynamic inhibition model suggests that activation of different populations of voxels might underlie temporal pattern separation. Thus, we used the additional similarity measures we used above (normalized dot product and norm difference) to strengthen the empirical evidence that different populations of voxels, rather than changes in the level of activation (potentially in a subgroup of voxels), underlies the results observed in DG. As above, if the pattern similarity results reflect changes in the activation of distinct voxels and not in the activation level of the same voxels, we predicted that we would replicate our correlation results in the normalized dot product measure, but not in norm differences. Indeed, mean normalized dot product values in the left DG showed a similar pattern to Pearson’s correlation: lower normalized dot product for items that are close in time, only within event, but not across events (Figure 5). As in Pearson’s correlation, in the 5^th^ Repetition, the interaction between Event and Temporal Distance was significant (*χ*^2^ = 6.77, *p* = .009, AIC reduction: 4.77). Temporal Distance significantly explained variance within event, but not across events (within event: *χ*^2^(1) = 10.429, *p* = .001, AIC reduction: 8.43; across events: *χ*^2^ = .33, *p* > .56). Critically, we found no difference based on temporal distance or events, or their interaction in the norm difference measure (all p’s > .14). Together, the results from the normalized dot product and the norm difference measures provide further evidence that temporally close items within the same event are represented by different populations of voxels in the left DG. These results are consistent with the dynamic inhibition model.

### CA3-DG connectivity increases with learning

Because we see that DG temporal differentiation is sensitive to event structure, it remains an open question how DG is sensitive to the event structure. Our results also show that over learning, neural patterns in CA3 become more stable within events. Thus, one hypothesis is that feedback or communication between CA3 and DG may also increase as learning advances. Indeed, a prominent theoretical model proposes that inhibition in DG, and consequentially pattern separation, is achieved via back-projections from CA3 (Myers & Scharfman, 2011; Scharfman, 2007). If pattern separation is mediated via CA3 back-projections and was higher in repetition 5 compared to early in learning, we aimed to examine if a concomitant increase in CA3-DG synchronization is evident from the first to the 5^th^ learning repetition. To address that, we used a background connectivity approach. Background connectivity is the correlation between slow fluctuations of BOLD signal in two ROIs, after removing trial-evoked activity using a linear regression (here we used 01-.035 Hz, slower than the rate of object presentation). It is thought to capture broad, state-level, changes in circuit dynamics (rather than momentary fluctuations), and has been shown to capture different phases of learning in hippocampal subfields (Duncan et al., 2014; Tompary, Duncan, & Davachi, 2015). We found that background connectivity between CA3 and DG did indeed significantly increase from the first to the 5^th^ repetition (1^st^ rep: *M* = .34, *SD* = .24; 5^th^ rep: *M* = .49, *SD* = .23, *t*(29) = 3.07, *p* = .005, Cohen’s *d* = .56; see Supplementary Figure 9 for data from all repetitions).

### No event representation in CA1

Although we focused on CA3 vs. DG representations during learning of events, for comparison purposes, we report data from CA1 as well. In the left CA1, we found no effect of Event or Temporal distance, nor an interaction between these effects, or any interaction with Repetition (*χ*^2^ ‘s < 2.21, *p*‘s > .13). In the right CA1, we found an interaction of Event by Temporal Distance (*χ*^2^ = 4.24, *p* = .039, AIC reduction: 2.24), with no Event by Temporal Distance by Repetition interaction (*χ*^2^ = 1.40, *p* = .24). Further, no main effects of Event and Temporal Distance, nor their interaction with Repetition, were significant (*χ*^2^ ‘s < 1.73, *p*‘s > .18). This is consistent with previous human studies that did not find CA1 multivariate pattern representations of spatial context (Stokes et al., 2015) or temporal context (Dimsdale-Zucker et al., 2022).

## Discussion

Decades of research have shown that the brain segments continuous experience into discrete events based on context (Clewett & Davachi, 2017; DuBrow & Davachi, 2014; Ezzyat & Davachi, 2011; Gernsbacher, 1985; Heusser et al., 2018; Radvansky & Zacks, 2017; Schank & Abelson, 1977; Zacks, 2020; Zacks et al., 2007). The hippocampus is central to event cognition (Ben-Yakov & Henson, 2018; Clewett et al., 2019; Maurer & Nadel, 2021; J. Zheng et al., 2022), potentially through its role in representing contextual and sequential information (Bellmund et al., 2020; Buzsáki & Moser, 2013; Buzsáki & Tingley, 2018; Davachi & DuBrow, 2015; Deuker et al., 2016; Eichenbaum, 2017; O’Keefe & Nadel, 1978; Ranganath & Hsieh, 2016; Rueckemann et al., 2021; Umbach et al., 2020). Critically, the hippocampus does not merely create copies of the external environment. Rather, it creates integrated representations of related life events, potentially allowing generalization and inference, while at the same time facilitating separated neural representations to distinguish memories (Brunec et al., 2020; Duncan & Schlichting, 2018; Liu et al., 2022; Molitor et al., 2021; Sugar & Moser, 2019). Theory and empirical data have broadly implicated the CA3 subregion of the hippocampus in integration and the DG subregion in separation (Knierim & Neunuebel, 2016; Marr, 1971; O’Reilly & McClelland, 1994; Treves & Rolls, 1994; Yassa & Stark, 2011). However, how time and context shape integration versus separation in hippocampal subregions to structure event representations, and how these representations change with learning, is unknown.

Here, participants learned repeating events while undergoing high-resolution fMRI. Event structure was evident in participants’ behavior, reflected in slower RTs for boundary items during learning, and slower RTs in a temporal memory test when memories bridged across different events. Using high-resolution fMRI and multivoxel pattern similarity analysis, we show novel evidence that CA3 activity patterns within the same event became more similar compared to across events, irrespective of temporal distance, suggesting that CA3 clustered representations based on events. The strength of neural event clustering in CA3 correlated with faster RTs during event transitions, suggesting that CA3 clustering facilitated transitioning between events. This adaptive CA3 segmentation is consistent with an attractor dynamics account for CA3.

In contrast to CA3, DG showed greater separation between objects that were close in time, compared to further in time. Previous human and rodent work showed evidence for pattern separation of visual stimuli or similar spatial environments (Baker et al., 2016; Bakker et al., 2008; Berron et al., 2016; Danielson et al., 2016; Kirwan & Stark, 2007; J. K. Leutgeb et al., 2007; S. Leutgeb & Leutgeb, 2007; Nakazawa, 2017; Neunuebel & Knierim, 2014; Wanjia et al., 2021). Here, for the first time to our knowledge, we show that DG performs differentiation across time, for temporally proximal information during an event, to facilitate distinct representations of temporally adjacent subevents. Importantly, our findings suggest that DG temporal differentiation is adaptive: when subevents belonged to different higher-level events that could be disambiguated based on perceptual context – in our case, the different background colors – we saw no temporal differentiation. A dynamic inhibition model, proposing that activated units are temporarily inhibited and then gradually released from inhibition, accounted for DG temporal pattern separation. In both CA3 and DG, integrated and differentiated event representations strengthened with learning. Finally, we used additional similarity measures to provide supporting evidence that different populations of voxels contributed to changes in similarity in both regions, consistent with attractor dynamics in CA3 and dynamic inhibition in DG.

CA3 event representations emerged even though the task allowed integration across different contexts. Specifically, in the current study, the order of objects and hence sequential events was replicated in the five learning lists. This theoretically offered participants the opportunity to predict and, thus, sequentially link across events even with the change in context features. From one perspective, that event segmentation arises from a prediction error process (Franklin et al., 2020; Zacks, 2020; Zacks et al., 2007, 2011), we might have expected to see a reduction in segmentation over learning repetitions as any ‘error’ signaling across events should be reduced with repeated exposure. Instead, behavioral markers of segmentation were still evident in the fifth repetition, and CA3 neural clustering was even stronger, suggesting that event segmentation is a process used and applied even when events are highly predictable. Consistent with this, previous studies have found evidence for hippocampal event representations in highly predictable environments (Hindy et al., 2016; Hsieh et al., 2014; Kok & Turk-Browne, 2018; Kyle, Stokes, et al., 2015; Schapiro et al., 2012, 2016). But, they used paradigms in which the target sequence was determined by high predictability compared to low predictability between sequences, thus learning the predictability structure was the only way of segmenting sequences into events (Hsieh et al., 2014; Schapiro et al., 2012, 2016). Other studies trained participants on discrete sequences or environments, requiring no segmentation (Dimsdale-Zucker et al., 2022; Hindy et al., 2016; Kok & Turk-Browne, 2018; Kyle, Stokes, et al., 2015; Liu et al., 2022; L. Zheng et al., 2021). By contrast, in our study, the fixed order made transitions within and across events fully and equally predictable, such that participants could learn and integrate across events. Nevertheless, participants leveraged the predictability in the list structure to adaptively chunk events instead of integrating across event boundaries (Clewett & Davachi, 2017; Shin & DuBrow, 2021).

In contrast to CA3 event-level representations, DG differentiated the representations of items that were close in time and experienced in the same event. Thus, DG temporal differentiation is specific and adaptive: when occurrences could be disambiguated based on different contexts, we did not observe separation in time. These results build on prior theories of the hippocampus (Kesner, 2018; Kesner & Rolls, 2015; Treves & Rolls, 1994; Yassa & Stark, 2011) as well as empirical work in humans showing reduced hippocampal pattern similarity between the multivoxel activity patterns of stimuli that are visually highly overlapping (Chanales et al., 2017; Favila et al., 2016; Fernandez et al., 2023; Koolschijn et al., 2019; Schlichting et al., 2015; Wanjia et al., 2021) or close in narrated, but not actual, time (Bellmund et al., 2022; but see Deuker et al., 2016). Additional studies have reported decreased similarity in CA3/DG between objects that share spatiotemporal context (Copara et al., 2014; Dimsdale-Zucker et al., 2018; Kyle, Smuda, et al., 2015; Liu et al., 2022), and specifically in DG (Berron et al., 2016; see also Baker et al., 2016). Our results extend this prior work in two critical ways: first, we show that DG differentiation can occur in the *temporal domain*, thus advancing our knowledge of how temporally extended experiences are represented in the brain (Davachi & DuBrow, 2015; Eichenbaum, 2014). Second, our results suggest that temporal differentiation is sensitive to context: DG does not separate all temporally similar experiences, but only when these experiences share the same context. Like the CA3 findings suggesting integration of temporally extended events, the DG temporal pattern separation results suggest that the hippocampus edits events, creating integrated and separated representations that are putatively adaptive for behavior (Ben-Yakov & Henson, 2018; Clewett et al., 2019; Sugar & Moser, 2019).

Motivated by the neurophysiological literature showing uniquely high levels of inhibition in DG (Bartos et al., 2007; Chawla et al., 2005; Coulter & Carlson, 2007; Freund & Buzsáki, 1998; Hainmueller & Bartos, 2020; Jinde et al., 2013) we considered that inhibitory dynamics in DG may underlie temporal differentiation. Potentially, voxels that are activated in one item, are then inhibited in the following item, maximizing separation between items. Then, this inhibition gradually decays, resulting in gradually decaying separation (Supplementary Figure 4). This is consistent with a model that proposes that back-projections from CA3 inhibit DG neurons, and facilitate pattern separation (Myers & Scharfman, 2011). Here, we found that CA3-DG connectivity increased with learning, concomitant to the increase in DG temporal pattern separation, providing some initial support to the later account. Future research could investigate the specific mechanism by which temporal pattern separation may arise in the DG.

An interesting question is whether hippocampal representations extend to higher, as well as lower, levels of event hierarchy. In our study, an even higher-level event could be the list, encompassing a sequence of color-defined events. The current study was not designed to test hippocampal representations at the list level, as each list (and repetition) was included in a separate fMRI scan. Thus, comparing representations within versus across lists would include comparing similarity values within vs. across scans, which is problematic (Mumford et al., 2014). Another related question is to what extent the breaks we imposed between repetitions of the same list influenced learning of hierarchical event representations in the hippocampus. Previous studies mentioned above showed event representations of discrete sequences, including at the list level (Dimsdale-Zucker et al., 2022; Hindy et al., 2016; Kok & Turk-Browne, 2018; Kyle, Stokes, et al., 2015; Liu et al., 2022; L. Zheng et al., 2021). Other studies showed hierarchical representations across multiple levels of hierarchy in the hippocampus (McKenzie et al., 2014; Theves et al., 2021), as well as time and space representations across extended periods of time (Hainmueller & Bartos, 2018; Nielson et al., 2015; Ziv et al., 2013). Thus, we postulate that the CA3 findings reported here could extend to higher hierarchical levels, whereby the representations are more similar within higher-level events that span a longer time (e.g., list) and might even be discontinuous.

In contrast, another theoretical possibility is an all-or-none event representation, whereby any change of features marks a novel event, and different levels are all the same in CA3. We believe that this possibility is less likely, given that in a lower event level, that of single items, changes of features between items did not interrupt CA3 clustered representations (we defined here items as subevents, but any subevent is also an event, and indeed vast memory literature defines each occurrence of an item as an ‘event’ to be remembered). It would be interesting to test, potentially using electrophysiological methods that have the appropriate temporal resolution, whether the same CA3 and DG dynamics we observed here at the event level also occur within each item in a fine-grained temporal scale.”

Many previous representational similarity studies in humans collapsed across CA3 and DG (e.g., Dimsdale-Zucker et al., 2018, 2022; Hindy et al., 2016; Kyle, Stokes, et al., 2015; Liu et al., 2022; Schapiro et al., 2012; Stokes et al., 2015; Wanjia et al., 2021; L. Zheng et al., 2021), hindering the examination of differences in these regions’ putative functions, namely CA3 integration through attractor dynamics compared to DG pattern separation (Kesner & Rolls, 2015; Marr, 1971; Treves & Rolls, 1994; Yassa & Stark, 2011). Here, we leveraged a widely accepted automatic segmentation protocol of hippocampal subfields (Iglesias et al., 2015), in combination with automated alignment of the ROIs from anatomical to functional space (Methods). In our fMRI data, we were able to dissociate CA3 and DG in two different measures: average BOLD activity and pattern similarity. Thus, while we do not wish to argue that we have achieved perfect anatomical segmentation, our segmentation procedure was useful to uncover compelling functional distinctions. Future studies, for example using 7T fMRI, will be important to confirm these findings. However, our novel procedure has clear advantages: it is available to any 3T scanner user, and it is fully automated, which is efficient and reproducible.

We did not find robust event or temporal representations in CA1. In this, we are consistent with previous human studies that did not find CA1 multivariate pattern representations of spatial context (Stokes et al., 2015) or temporal context (Dimsdale-Zucker et al., 2022). Shared by the current and these previous studies is that context representations were examined with only minimal learning of a few repetitions. Interestingly, other human studies that used extensive learning show spatial and temporal context representation in CA1 (Hindy et al., 2016; Kyle et al., 2015; Schapiro et al., 2012; Thavabalasingam et al., 2019). In rodents as well, some spatial and temporal representations in CA1 requires learning (Gill et al., 2011; Lever et al., 2002; Mankin et al., 2012; Pastalkova et al., 2008). It is possible that more extensive learning is required for context representations to manifest in CA1, because of CA1’s diverse inputs, namely CA3 and entorhinal cortex (Amaral, 1993; Kesner & Rolls, 2015; Knierim, 2015; Marr, 1971). Theoretical models and emerging empirical work converge on the notion that in familiar environments, CA1 activity is more strongly modulated by CA3 inputs, whereas in novel environments, entorhinal input has a strongly modulatory effect on CA1 (Bein et al., 2020; Colgin, 2016; Colgin et al., 2009; Duncan et al., 2014; Hasselmo et al., 1996, 2002; Kemere et al., 2013; Lopes-dos-Santos et al., 2018; Tort et al., 2009). Potentially, when an environment is very well-learned, CA3 input to CA1 is strong enough such that CA3 contextual representations also shape CA1 representations.

In sum, in the present study we examined learning of temporally extended events and found that CA3 integrated items within event, and separated across events, consistent with an attractor dynamics account. DG, in contrast, separated representations for items that were close in time and within the same context, consistent with temporal differentiation that is sensitive to context. These results show how the hippocampus hierarchically represents events to address a major challenge of our memory system: how to integrate information to build contextually relevant knowledge, while maintaining distinct representations of this same information (Brunec et al., 2020; Duncan & Schlichting, 2018; McKenzie et al., 2014; Tompary & Davachi, 2017).

Through learning, such hierarchical event representations can become the building blocks of organized semantic representations, including schemas, categories, and concepts (Collins & Quillian, 1969; Eichenbaum, 2004; Garvert et al., 2017; Ghosh & Gilboa, 2014; Mack et al., 2018; Morton et al., 2017; Murphy, 2004; Nieh et al., 2021; Renoult et al., 2019; Theves et al., 2021; Tulving, 1972). How event representations as observed here are transformed to become schematic knowledge is an exciting question for future research (Gilboa & Moscovitch, 2021; Moscovitch et al., 2016; Robin & Moscovitch, 2017; Tompary & Davachi, 2017).

## Author contributions

O.B. and L.D. conceptualized the experimental design and analytical approach. O.B. collected and analyzed the data. O.B. wrote an initial draft of the paper, O.B. and L.D. revised and edited the paper. L.D. supervised and obtained funding.

## Data and code availability

Data will be available upon a reasonable request from the authors. The code will be available upon publication.

## Acknowledgments

This research was supported by the National Institute of Mental Health Grant R01MH074692 toL.D. O.B. was supported by the National Institute of Mental Health Grant T32MH065214, the McCracken fellowship, and a Grinker award. We thank Andrew Heusser and J. Quinn Lee for their help in the initial conceptualization of the behavioral paradigm, and Natalie Plotkin and Adam Benzekri for their help in behavioral piloting of the task. We thank Monika Riegel, Tarek Amer, and David Clewett for insightful comments and conversations, and Emily Cowan, Tarek Amer, Camille Gasser, and John Thorp for comments on earlier drafts of the manuscript.

The authors declare no conflict of interest.

## Footnotes

1 Future analyses of this mutli-task design include analyses of temporal memory, color memory, and pre-post similarity from the random-order viewing, as well as analysis of cortical areas in the list-learning task.

2 Between the last repartition of each list and the temporal memory participants observed all items in random order. We expected that given the extensive training that included many repetitions, and since we presented all items regardless of events, this will have minimal influence on behavior. Indeed, we observed high accuracy rates and confidence levels.

