## Supplemental Information for "Event integration and temporal differentiation: how hierarchical knowledge emerges in hippocampal subfields through learning"

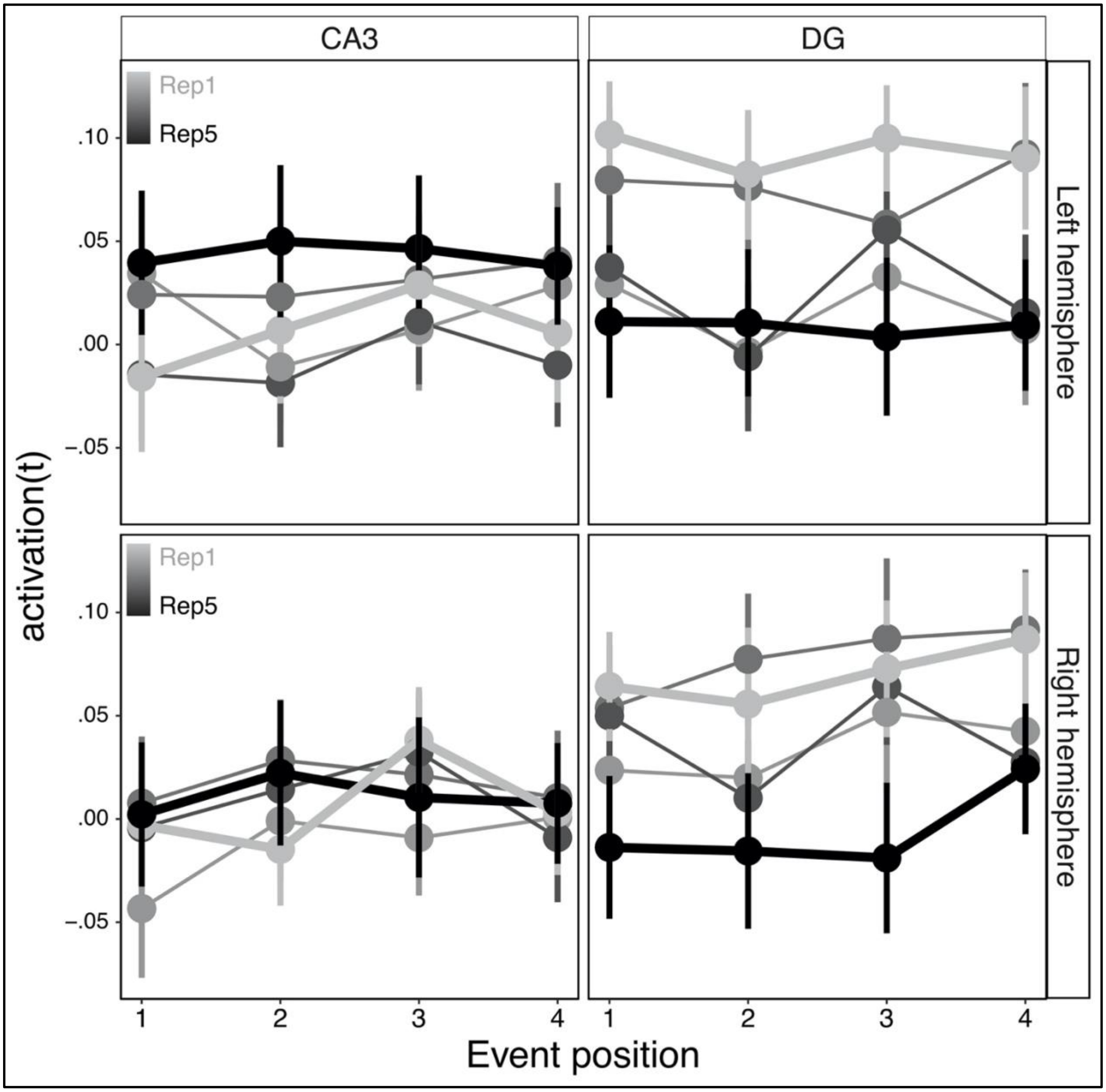


*Supplementary Figure 1.* Univariate activation (average t-statistic) per each ROI (CA3/DG) and in each hemisphere (left/right). Rep1: 1st repetition. Rep5: 5th repetition. Data are presented as mean values, error bars reflect +/- SEM. N = 30. For details and statistical analysis, see main text.

| ROI/Repetition | 1 | 2 | 3 | 4 | 5 |
|---|---|---|---|---|---|
| CA3 | .006(.13) | .001(.14) | .023(.15) | .000(.13) | .027(.16) |
| DG | .082(14) | .025(.15) | .077(.17) | .03(.15) | .001(.17) |

*Supplementary Table 1.* Mean univariate activation (average t-statistic) per repetition and ROI (CA3/DG) averaged across hemisphere and event position (as no effect of hemisphere or position was found, see main text and *Supplementary Figure 1*). SD are in parentheses.

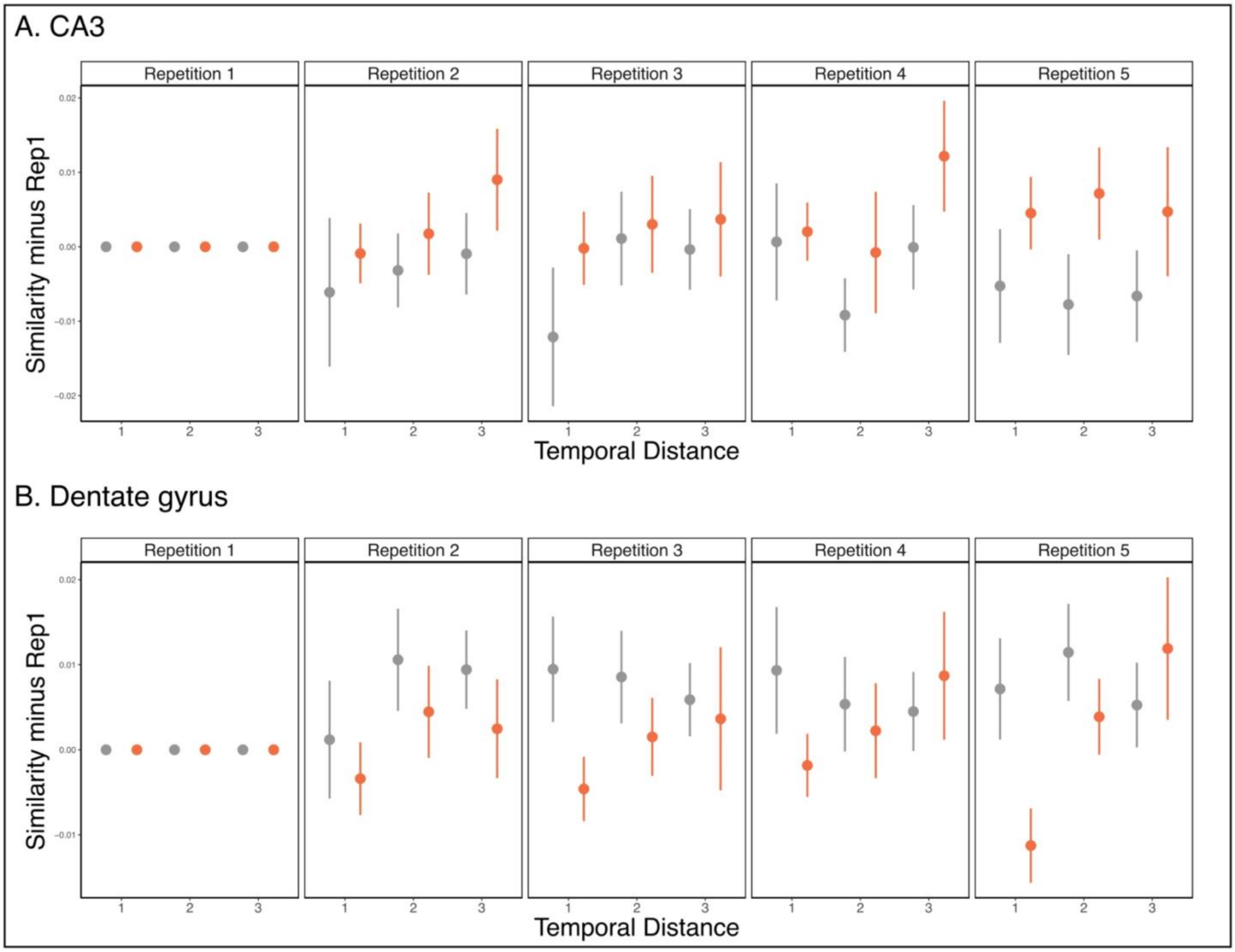


*Supplementary Figure 2.* Similarity values in all repetitions, in A. CA3, and B. Dentate gyrus (DG). Orange: similarity within-event. Grey: similarity across events. The similarity values of the first repetition (Rep1) are subtracted from all repetitions. Data are presented as mean values, error bars reflect +/- SEM. N = 30. For details and statistical analysis, see main text.

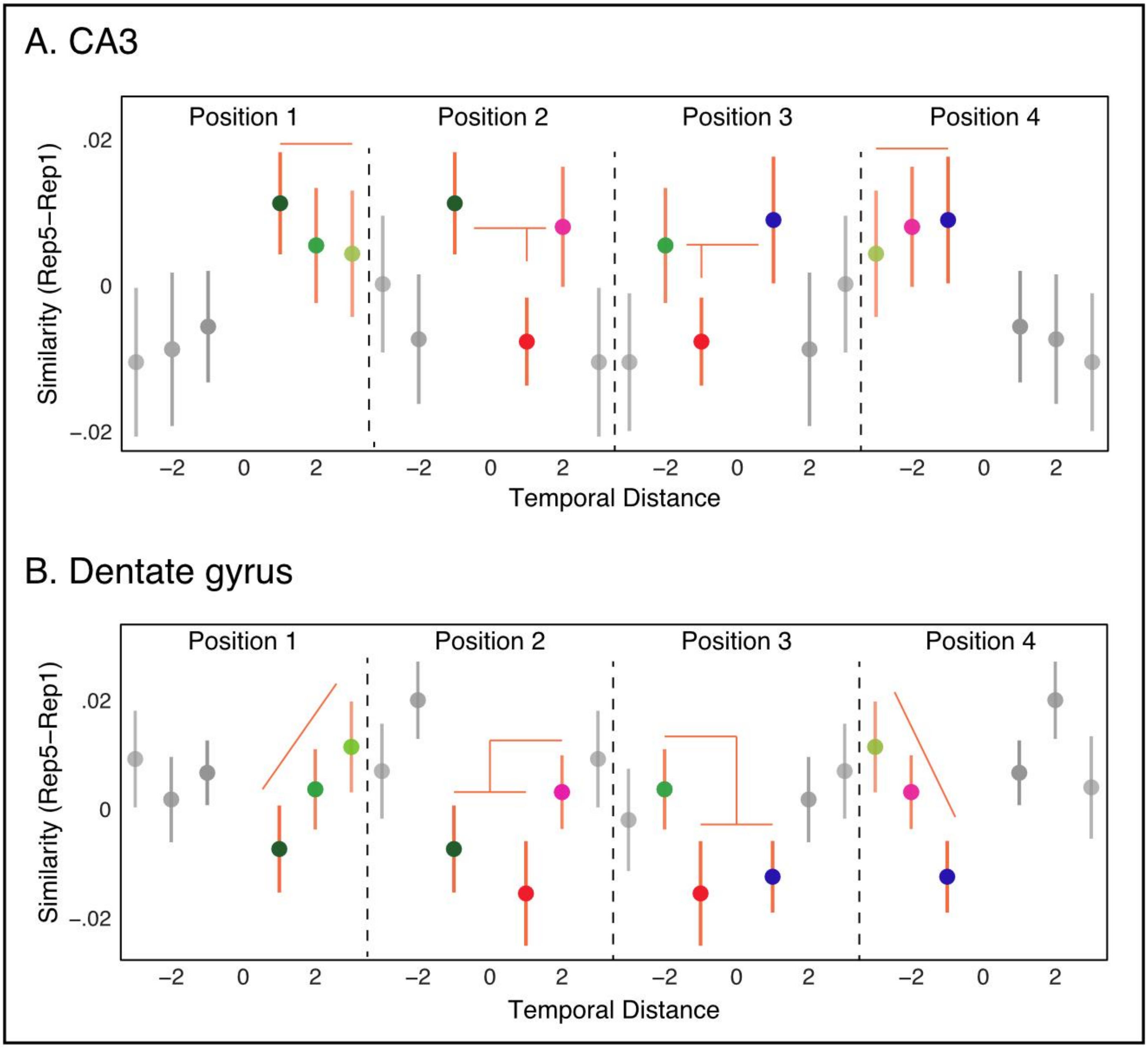


*Supplementary Figure 3*. Representational similarity per event positions in A. CA3, and B. Dentate gyrus (DG). The analyses in the main text collapsed across all positions (1-4) within an event. As an exploratory analysis, in both CA3 and DG, we examined the stability of representational similarity effects across event positions. In the 5$^{th}$ repetition, it seems like in CA3 similarity with the edges of the event, namely event positions 1 and 4 was higher than similarity in the middle of the event, between positions 2 and 3, was lower. In DG temporal pattern separation (Fig. 6, main text) was stable throughout the event. That is consistent with our dynamic inhibition model (Fig. 7, main text). Orange: similarity within-event. Grey: similarity across events. Note that each data is mean similarity between two event positions. For visualization, each data point is presented twice, each time anchored based on the panel per each event position. Same data points are represented using the same fill color. For example, the similarity between event position 1 and 4 is presented in light green, once in the panel anchoring on position 1, and once in the panel anchoring on position 4. The similarity values of the first repetition (Rep1) are subtracted from all repetitions. Data are presented as mean values, error bars reflect +/- SEM. N = 30.

### Control for univariate activation

Note that the Pearson's correlation measure subtract the mean, and therefor accounts for differences in mean activation. Nevertheless, to further establish that differences in representational similarity cannot be attributed to differences in univariate activation, we repeated the main analyses, but adding univariate activation as an explaining variable to the statistical models. Specifically, we added the difference in univariate activation between the fifth and the first repetition. Since each similarity value is the correlation between two items, we took the univariate activation difference per each trial, for each of the corresponding two items in the similarity analysis, as well as the interaction between them. This made a total of three univariate activation explaining variables added to our mixed-level linear models: item1, item2, item1*item2. Shortly, the results reported in the paper did not change when including univariate activation in our statistical models, indicating that our similarity results are unlikely to be attributed to changes in univariate activation. We report these analyses here in detail.

In the left CA3, we found a main effect of Event, which remained significant also when controlling for univariate activation (within vs. across: $\chi^2_{(1)} = 8.97$, $p = .0028$, AIC diff.: 6.97). The interaction of Event by Repetition (1-5) remained marginally significant ($\chi^2_{(1)} = 3.61$, $p = .057$; AIC diff.: 1.61). There was no main effect of Temporal Distance ($\chi^2_{(1)} = 2.22$, $p = .13$), nor interaction effects for Event by Temporal Distance ($\chi^2_{(1)} = .04$, $p = .84$) or of Event by Temporal Distance by Repetition ($\chi^2_{(1)} = .01$, $p = .92$). In the 5$^{th}$ repetition, within event similarity was significantly higher than across events similarity, controlling for univariate activation ($\chi^2_{(1)} = 5.31$, $p = .02$, AIC diff.: 3.31).

In the left DG, the interaction between Event and Temporal Distance was significant also when controlling for univariate activation ($\chi^2_{(1)} = 6.30$, $p = .01$, AIC diff.: 4.30), as was the effect of Temporal Distance within event ($\chi^2_{(1)} = 11.09$, $p = .0009$, AIC diff.: 9.09). The 3-way interaction of Event by Temporal Distance by Repetition remained significant as well ($\chi^2_{(1)} = 4.68$, $p = .03$, AIC diff.: 2.68). In the 5$^{th}$ repetition, there was a significant interaction of Event by Temporal Distance ($\chi^2_{(1)} = 6.22$, $p = .01$, AIC diff.: 4.22), stemming from lower pattern similarity for objects close vs. far in time within event (within event, main effect of Temporal Distance: $\chi^2_{(1)} = 10.12$, $p$

= .001, AIC diff.: 8.12; distance of 1 vs. 2: $\chi^2_{(1)}$ = 5.55, $p$ = .02, AIC diff.: 3.55; distance of 1 vs. 3: $\chi^2_{(1)}$ = 8.06, $p$ = .005, AIC diff.: 6.06). These results, in both the left CA3 and DG, did not change if we used raw activation values, rather than the difference from the first repetition.

**Schematic of similarity measures simulations**

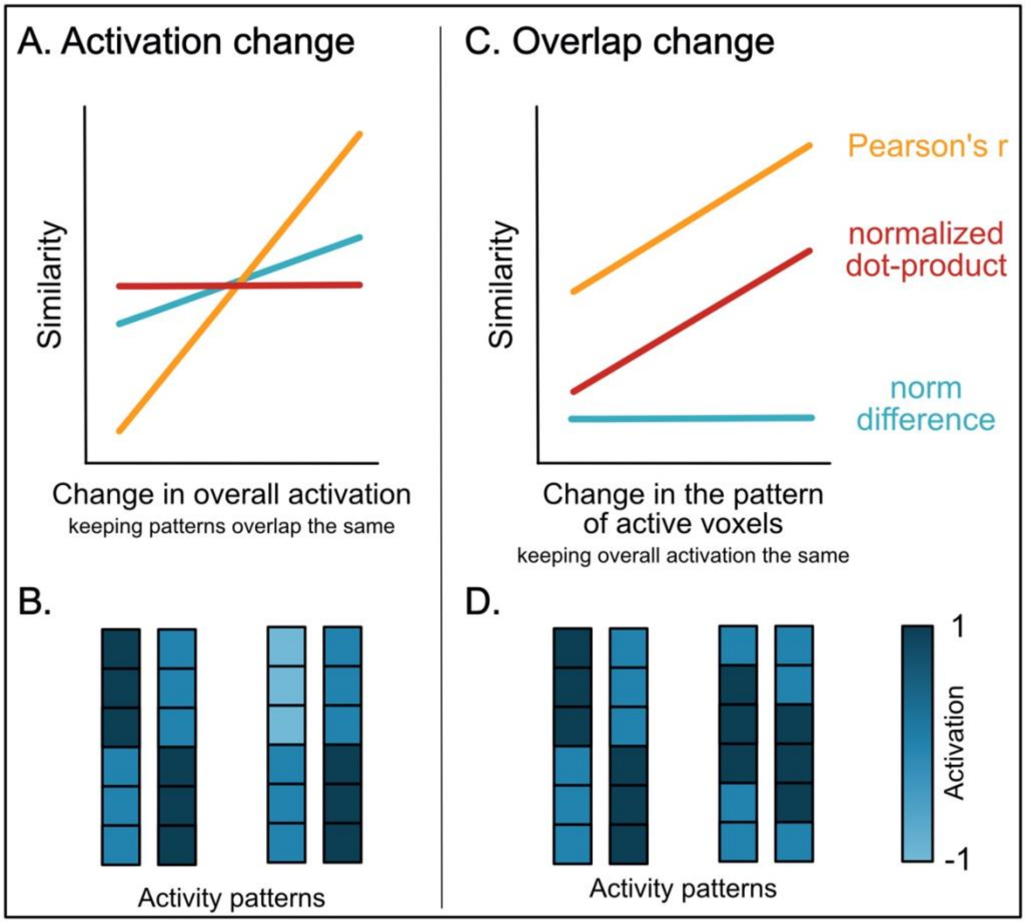


*Supplementary Figure 4. Schematic of the simulations of Pearson r, normalized dot product, and norm difference measures.* A. When active voxels do not overlap, changing the level of activation in a subgroup of voxels led to differences in Pearson's r and norm difference, but not in the normalized dot-product. B. Activity patterns showing the change in a subgroup of voxels. C. Altering the voxels' overlap, without changing their level of activation, led to a change in Pearson's r and normalized dot-product, but not in norm difference. D. Activity patterns showing the change in overlap of active voxels across both activity patterns. See more details in the main text in the Supplementary Information.

**Full simulation details: Pearson's R, normed dot product (ndp) and norm difference**

We assessed whether different populations of voxels, rather than changes in level of activation (potentially in a subgroup of neurons), may underly similarity differences. To that end, we adopted to human fMRI two known similarity measures that have been previously used in rodent work (Madar et al., 2019). These are the normalized dot product (ndp) and the vector-norm difference (see equations 1 and 2 below, X and Y refer to two activity patterns). The ndp is the sum of the product of each pair of parallel voxels in the two vectors (i.e., dot product), normalized by the norms of the two vectors (the norm of each vector is the dot product of the vector by itself, which gives the total length of the vector). The ndp can be thought of as the cosine of the angle between two vectors (i.e., higher ndp means more similar patterns). The vector-length difference is the difference between the norms of two activity patterns. Note that since each voxel is multiplied by itself, negative activation values will become positive and contribute to a larger norm of a vector. Thus, this measure is not an average activation level, but rather the sum of magnitude of changes in activation level, regardless of the direction.

$$(1)\ NDP: \frac{\sum_{i=1}^{N} X_i Y_i}{\sqrt{\sum_{i=1}^{N} X_i^2}\sqrt{\sum_{i=1}^{N} Y_i^2}}$$

$$(2)\ Norm\ Difference: \left|\sqrt{\sum_{i=1}^{N} X_i^2} - \sqrt{\sum_{i=1}^{N} Y_i^2}\right|$$

To provide intuition, we ran simulations demonstrating where the ndp, correlation, and norm-difference measures provide similar or divergent results. We ran this simulation for few different levels of activation. For all simulations, number of voxels in the patterns did not change the results, thus here we only report the results with 40 voxels (on average, participants had 39.41 voxels in the left CA3, range 21-68, and 47.8 voxels in the left DG, range 30-68).

First, we simulated a scenario whereby in one pattern half of a pattern is activated (e.g. [0,0,0,0,1,1,1,1]), whereas in the second pattern the other half is activated and changes its level of activation (e.g., [x,x,x,x,0,0,0,0], where x denotes a changing level of activation). As can be seen in Supplementary Figure 5A, this has no effect on the ndp. Because the population of voxels do not overlap, the ndp is always 0, no matter if activity levels changed in these non-overlapping voxels. However, these changes in activation level do influence the Pearson correlation: when x is negative, the patterns are positively correlated because in both patterns the first half is lower than the mean, whereas the second half is higher than the mean (and to the same extant since the correlation computes distance from the mean per sample in units of standard deviations). When x is positive, the patterns flip to be negatively correlated, because in one the first half is lower than the mean, whereas the second is higher than the mean, and in the second pattern, it is the exact opposite. It can also be seen that the norm-difference shows the expected result of larger differences the more the second pattern’s activation level is farther from 1 or -1. The symmetry around 1 and -1 is because the norm-difference is irrespective of the direction of change. Thus, this simulation demonstrates how the Pearson correlation is sensitive to changes in the mean activation of sub-part of the pattern, whereas the ndp is less so.

Next, we simulated a situation akin to partial repetition suppression, namely a well-known phenomena of reduced activation following repeated exposure (Grill-Spector et al., 2006). For example, if through learning, voxels come to represent two neighboring items, then upon their sequential presentation, voxels might show lower activation levels for the second item. We simulated this by having one pattern fully activated (e.g., [1,1,1,1,1,1,1,1]), whereas in the other we varied the percentage of non-active voxels (e.g., from [0,1,1,1,1,1,1,1,] to [0,0,0,0,0,0,0,1]). As can be seen in Supplementary Figure 5B, that led to expected increased norm-difference. The ndp measure decreases as the number of overlapping voxels decreases: with less activated voxels, it is as if there are less voxels 'pointing in the same direction' and thus the ndp decreases. The correlation measure stays roughly similar, because the relative distance of each point from the mean is not changing much in this simulation.

Finally, we simulated a situation whereby the activation levels and the number of activated voxels do not change, but the percentage of overlapping voxels does change. (e.g, one pattern was [0,0,0,0,1,1,1,1] and the other ranging from [x,x,x,x,0,0,0,0], which has no overlap to full overlap [0,0,0,0,x,x,x,x]). As expected, this influenced the ndp and the Pearson correlation, but did not influence the norm difference. As can be seen in Supplementary Figure 5C,D, more overlapping voxels increased the Pearson correlation and the ndp when activation values were positive, because that means that more voxels are pointing towards the same "direction" in the voxel space as our reference pattern, which had positive values, and the angle between the voxels get smaller. Naturally, when activation values are negative, namely, the voxels are pointing in the opposite direction in the voxel space compared to the reference pattern, then the more overlap there is, the lower the correlation and ndp value are. Together, these simulations show that the ndp measure is less sensitive to changes in levels of activation, compared to the Pearson correlation, but that it is sensitive to changes in the population of activated voxels. The norm difference is only sensitive to total amount of changes in activation, but not to changes in the voxels' population.

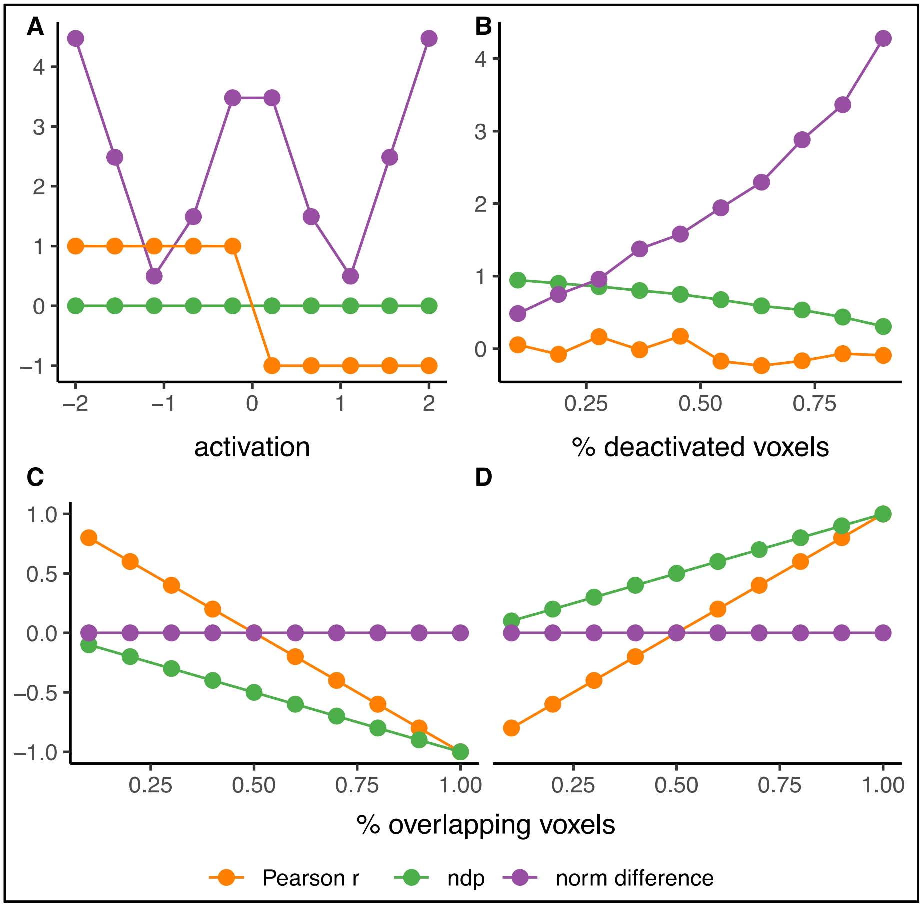


*Supplementary Figure 5. Simulations of Pearson r (orange), ndp (green) and norm difference (purple) measures.* A. maintaining voxels' overlap, while altering the level of activation. B. deactivating voxels in the activity pattern. C, D. altering the percentage of voxels' overlap, under two different activation levels. See main text for more details.

*A dynamic inhibition model for temporal differentiation in DG*

We formalized inhibitory dynamics to account for the lower similarity between temporally proximal items. Specifically, we propose the DG neurons that are activated for one item (e.g., item n) become inhibited for some time such that they are not active when the immediately following item (n+1) appears. This results in other neurons representing the n+1 item, and in a low overlap between activity patterns of temporally adjacent items (items n and n+1). Inhibition then gradually decays, so that for the following item (n+2), some of the neurons can overcome this inhibition and are active again, resulting in a slightly higher correlation between items n and n+2. By item n+3, inhibition further decays such that similarity levels are high again and reach the levels of between-event similarity. Between events, inhibition does not come into play because items can be distinguished based on perceptual features alone, thus DG differentiation is unnecessary. This idea can be shown conceptually in Supplementary Figure 6.

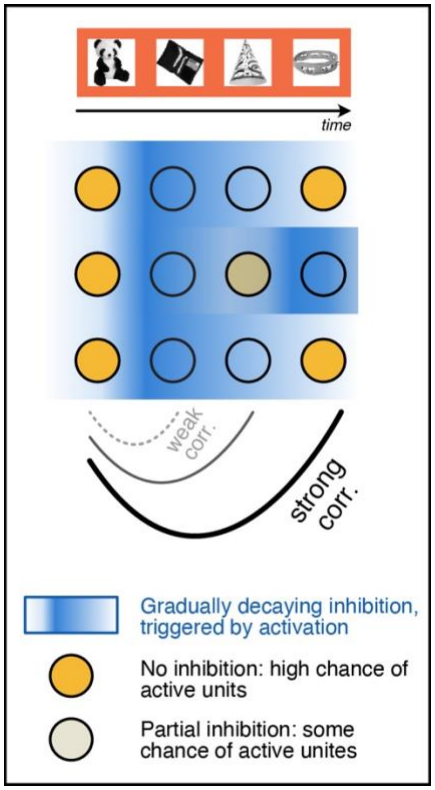


*Supplementary Figure 6. Dynamic Inhibition model for temporal differentiation.* Activated units (yellow circles) trigger inhibition that is maximal in the next trial (n+1) but then gradually decays (blue to white background). By trial n+2, partial inhibition leads to some chance for a unit to be activated again (faded yellow). By trial n+3 inhibition decays further, such that a unit has a high chance of being active. The curves reflect the resulting correlation between activity patterns.

*Details of the dynamic inhibition model and simulations*

Our primary goal in these simulations was to illustrate that the temporally decaying inhibition could, in principle, result in the pattern of the data we observed in the left DG. We do not aim to offer a comprehensive mechanistic account for temporal differentiation in DG (this would be beyond the scope of the current paper, but see Discussion, main text). We simulated activity patterns of 2 continuous events, to examine similarity values for all temporal distances within and across events. Patterns were set as vectors of 0 (inactive units) or 1 (active units). The portion of active units for each simulated activity pattern was set to .1 or .2, reflecting low levels of activation in DG (Chawla et al., 2005; Jung & McNaughton, 1993). Inhibition levels in the model determine the rate of units from a pattern if a previous item that are not allowed to be activated for a current pattern. Inhibition level linearly increased across repetitions, and exponentially decayed in time between items in each repetition, based on eq. (1):

$$(1)\ inhibition = (initial_{inhib} + (rep - 1) * increase_{inhib}) * \ e^{-\omega t}$$

The argument in the parentheses is the maximal level of inhibition in each repetition. $Initial_{inhib}$ is the level of inhibition in the first repetition (Rep1), reflecting minimal inhibition in DG. This should be larger than zero, capturing the idea that already in the first presentation, some level of inhibition is in place after neurons are activated. We arbitrarily set $\mathrm{Initial}_{\mathrm{inhib}}$ to .5. However, the result of changing this level of inhibition is trivial – it will be reflected in less inhibition during the initial stages of learning. $Increase_{inhib}$ sets a linear increase in each repetition. We chose a linear increase given the gradually increasing differentiation across repetition observed in the empirical data. We set $Increase_{inhib}$ to reach a level of 1 by repetition 5, reflecting full inhibition. Thus, the magnitude of the increase is fully derived from $\mathrm{Initial}_{\mathrm{inhib}}$. This resulted in $Increase_{inhib}$ = .125. As seen in eq. 1, inhibition decayed exponentially in time, using a scaling factor $\omega$. We explored a scaling factor of .15, .2, and .25. *t* denotes the temporal distance between items, and was set to Temporal Distance – 1, to have no decay for a Temporal Distance of 1 (i.e., between items n and n+1, t=0, $e^0 = 1$).

The simulation starts with the first presentation (Rep1), allocating a random activity pattern to the item at position 1 in the event. Then, for each of the following items within the event, a random activity pattern is generated, with inhibition levels determined based on eq. (1), setting the rate of units from the pattern of the previous item that is not allowed to be activated in the pattern of the current item. For each item, inhibition of units in previous patterns in the event is enforced. Precisely, for an item in position 2, units from position 1 are inhibited. For an item in position 3, units from positions 1 and 2 are inhibited, each at different levels based on eq. (1), and for an item in position 4, units from positions 1,2, and 3 are inhibited, each at different levels based on eq. (1). Then, item 1 of the next event is presented to the model and the process starts again, by allocating a random activity pattern to item 1. This effectively turned off inhibition between events.

For each of the next repetitions, the model would attempt to activate the pattern set for each item in the previous repetitions, reflecting memory reactivation (Danker & Anderson, 2010; Ritchey et al., 2013; Tompary et al., 2016). Because inhibition levels increase across repetitions, that could mean that some overlap that was permitted in previous repetitions, might be too high in the current repetition, and some of the previous units should now be inhibited (their activity set to 0). To implement that, from the overlapping units between the reactivated pattern and the pattern of the previous item being examined, we randomly sampled the minimal number of units that should be set to 0 to answer current levels of inhibition, and set these units to 0.

Next, we computed Pearson's correlation between simulated patterns, based on Temporal Distance and within and across events, as was done for the empirical data. Supplementary Figure

7 depicts the simulation results in Rep5 for .2 activated values, across a range of ROI sizes and decay rates (scaling factor $\omega$). As can be seen, the model produced similar results to our empirical data. Supplementary Figure 8 shows all repetitions for an ROI size of 50 (similar to the average number of voxels in the left DG in our data). Noticeable differentiation emerges in Rep3, consistent with empirical data. As expected, when inhibition decays quickly (in this simulation, $\omega = .25$), it means that already at a temporal distance of 2, there are rather low levels of inhibition, and the similarity between patterns is restored.

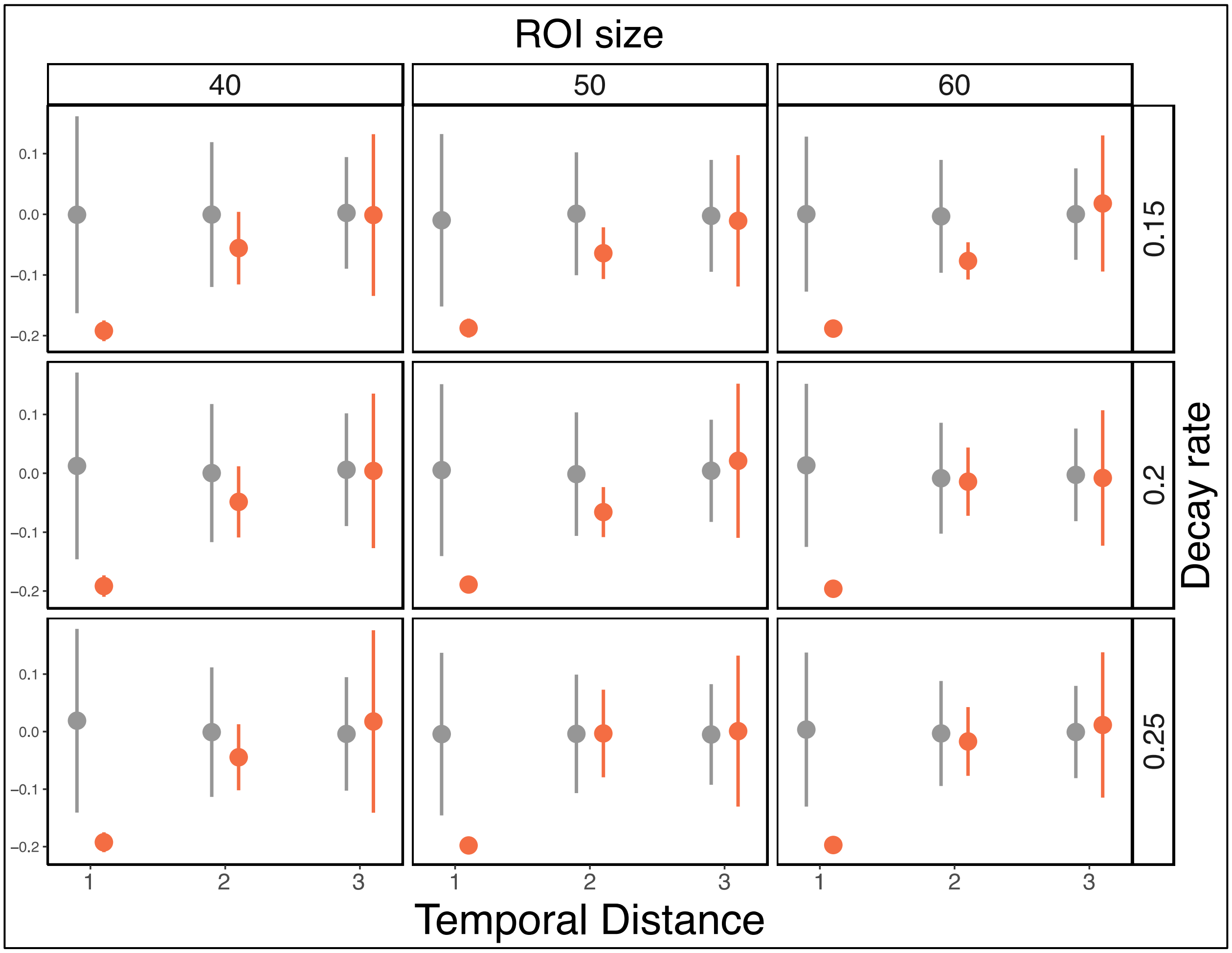


*Supplementary Figure 7.* Simulation results for the DG dynamic inhibition model, Rep 5. Similarity values for different temporal distances, within event in orange, and across events in grey. Data are presented as mean values over 500 simulations, error bars reflect +/- SD. For temporal distance of 1 there are no error bars, as the Pearson's correlation between patterns with no overlapping units is identical regardless of the specific units activated.

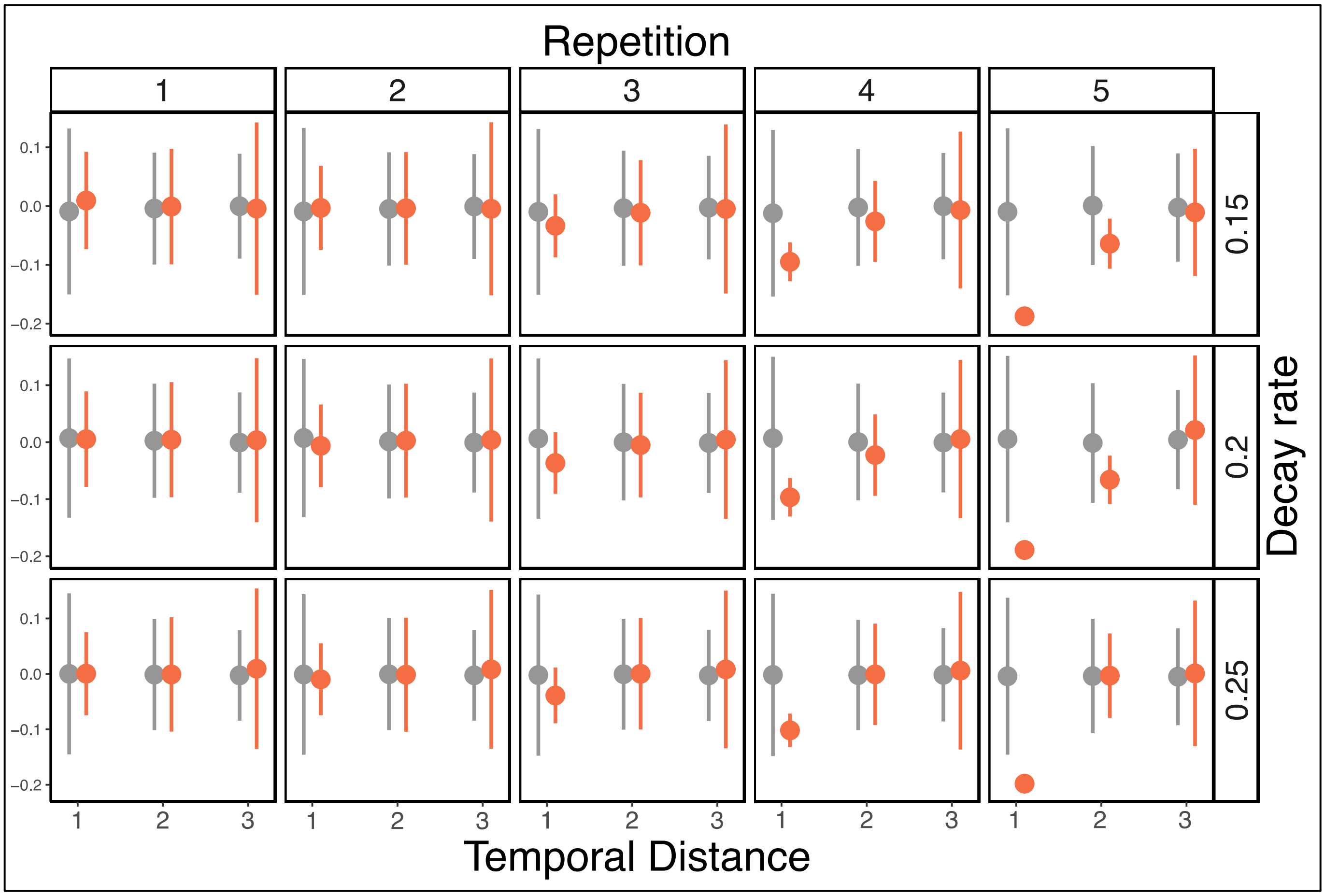


*Supplementary Figure 8.* Simulation results for the DG dynamic inhibition model for ROI size of 50 units. Similarity values for different temporal distances, within event in orange, and across events in grey. Data are presented as mean values over 500 simulations, error bars reflect +/- SD. As can be seen, already in Rep 3, some differentiation is observed. For temporal distance of 1 there are no error bars, as the Pearson's correlation between patterns with no overlapping units is identical regardless of the specific units activated.

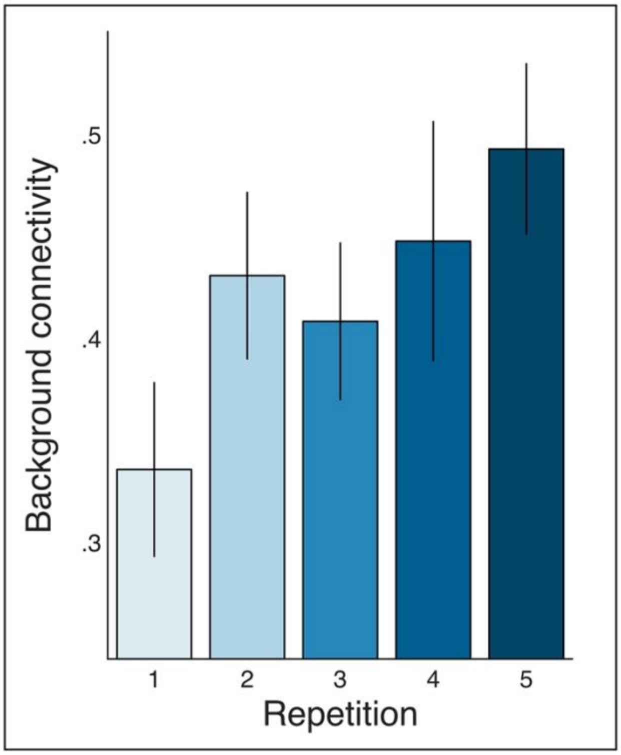


*Supplementary Figure 9.* Background connectivity between the left CA3 and DG in all repetitions. Data are presented as mean values, error bars reflect +/- SEM. N = 30. For details see main text.

Anterior versus posterior hippocampus

Although not the focus of our investigation, given the interest in the field (Brunec et al., 2018; Poppenk & Moscovitch, 2011; Thorp et al., 2022), we divided the hippocampus into thirds (Tompary and Davachi, 2017) and examined the anterior vs. posterior portions of the hippocampus. Since our primary hypotheses and results address different effects in DG versus CA3, and dividing the hippocampus into anterior/posterior mixes subfields, we did not a-priori expect any results when looking at multivariate similarity effects in the anterior or posterior hippocampus.

*Higher Univariate activation for boundaries in the posterior hippocampus*

The univariate activation estimates during learning (t-statistics resulting from a standard GLM analysis, as was done for CA3 and DG, see Methods: Univariate activation) per each event position and repetition and each participant, averaged across all voxels in each ROI, were entered into a repeated measures ANOVA with the factors of Event Position (1 through 4), Repetition (1 through 5), ROI (anterior/posterior), and Hemisphere (right/left). We found a main effect of Repetition ($F_{(1,29)} = 8.94$, $p = .006$, $\eta_p^2 = .24$), and of ROI ($F_{(1,29)} = 16.4$, $p = .0003$, $\eta_p^2 = .36$), but no interaction between Repetition and ROI ($F_{(1,29)} = .25$, $p = .62$). A marginal main effect of Hemisphere was observed as well ($F_{(1,29)} = 3.49$, $p = .072$, $\eta_p^2 = .11$), and a significant interaction of Hemisphere and Repetition ($F_{(1,29)} = 4.88$, $p = .035$, $\eta_p^2 = .14$). While no main effect of Event Position was found ($F_{(1,29)} = .002$, $p = .96$), there was a significant interaction of Event Position and ROI ($F_{(1,29)} = 11.19$, $p = .002$, $\eta_p^2 = .28$). All other effects were not significant ($F\text{'s}_{(1,29)} < 1.74$, $p\text{'s} > .19$). As can be seen in Supplementary Figure 10, this reflects higher activation of the boundary item (event position 1) in the posterior hippocampus, consistent with previous findings (Ben-Yakov, Eshel & Dudai, 2013; Ben-Yakov Robinson & Dudai, 2014). Given these prior findings, we followed up on the Event Position by ROI interaction, by looking directly at the boundary effect, namely univariate activation at event position 1 vs. all event positions 2,3,4, in each ROI. We collapsed across hemispheres since no interaction of Event Position by ROI by Hemisphere was found. Univariate activation in each ROI entered a repeated-measures ANOVA with the effect of Boundary (event position 1 vs. the average of event positions 2,3, and 4) and Repetition (1 through 5, as above). We found a significant effect of Boundary in the posterior hippocampus ($F_{(1,29)} = 11.27$, $p = .002$, $\eta_p^2 = .28$), and of Repetition ($F_{(1,29)} = 6.97$, $p = .013$, $\eta_p^2 = .19$), and no interaction ($F_{(1,29)} = 2.02$, $p = .17$). The same analysis in the anterior hippocampus yielded a significant effect of Repetition ($F_{(1,29)} = 5.9$, $p = .02$, $\eta_p^2 = .$), but no effect of Boundary nor an interaction ($F$'s$_{(1,29)}$ < .71, *p's* > .40). This is different from prior reports of a boundary response in both the anterior and posterior hippocampus (Ben-Yakov, Eshel & Dudai, 2013; Ben-Yakov Robinson & Dudai, 2014).

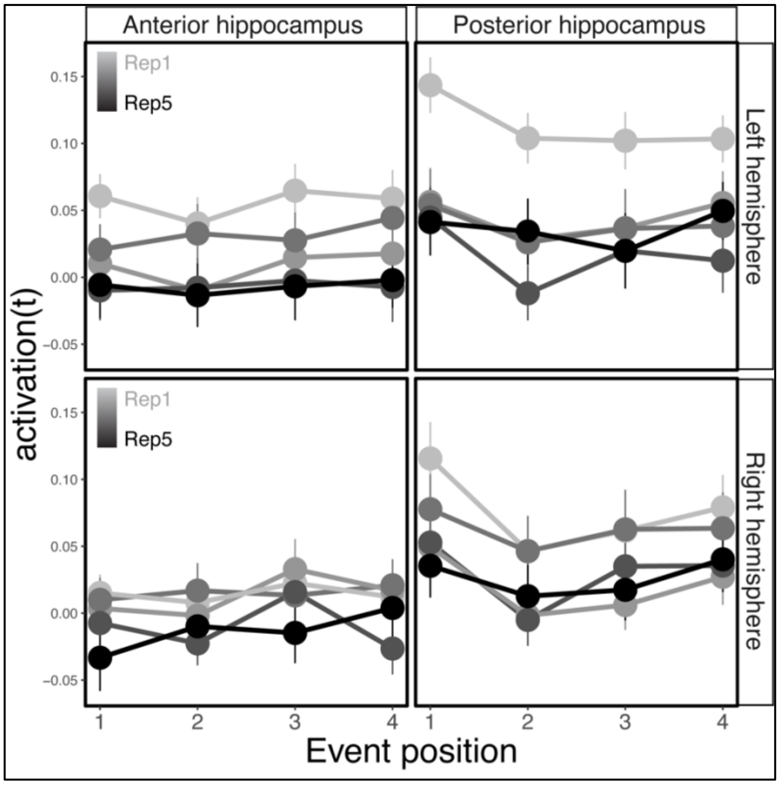


*Supplementary Figure 10.* Univariate activation (average t-statistic) per each ROI (anterior/posterior hippocampus) and in each hemisphere (left/right). Rep1: 1st repetition. Rep5: 5th repetition. Data are presented as mean values, error bars reflect +/- SEM. N = 30. For details and statistical analysis, see above.

*Item-item representational similarity analysis*

We took the same approach as with the CA3 and DG (see Methods). First, the similarity values per pair of objects in repetitions 2-5 were entered into a mixed-effects linear model that examined the interaction between Event (within event/across events), ROI (anterior/posterior), and Hemisphere (right/left), finding no interaction of Event by ROI by Hemisphere ($\chi^2_{(1)}$ = .0129, $p$ = .91). This analysis did reveal an interaction of Event by ROI ($\chi^2_{(1)}$ = 8.53, $p$ = .003, AIC reduction: 6.53), and a marginally significant interaction of Event by Hemisphere ($\chi^2_{(1)}$ = 3.51, $p$ = .061, AIC reduction: 1.61). Even though the 3-way interaction of Event by ROI by Hemisphere was not significant, given these 2-way interactions, we examined each ROI in each

Hemisphere separately for changes in event representations through learning. Within each of the left and right anterior and posterior hippocampal ROIs, we tested the effects of Event, Temporal Distance, and Repetition, and the interaction between these factors (see Methods). Generally, there was no effect of Repetition nor an interaction of Repetition in any ROI. Thus, Supplementary Figure 11 presents the data averaged across repetition.

Here are the significant and marginally significant results:

right anterior: Event ($\chi^2_{(1)} = 7.96$, $p = .005$, AIC reduction: 5.96), Event by Temporal Distance interaction ($\chi^2_{(1)} = 2.94$, $p = .087$, AIC reduction: 3.52).

right posterior: Temporal Distance ($\chi^2_{(1)} = 3.49$, $p = .061$, AIC reduction: 1.49), Event by Temporal Distance interaction ($\chi^2_{(1)} = 3.73$, $p = .053$, AIC reduction: 1.73).

No other effects were found in any ROI ($\chi^2$'s$_{(1)} < 2.2$, $p$'s > .14)."

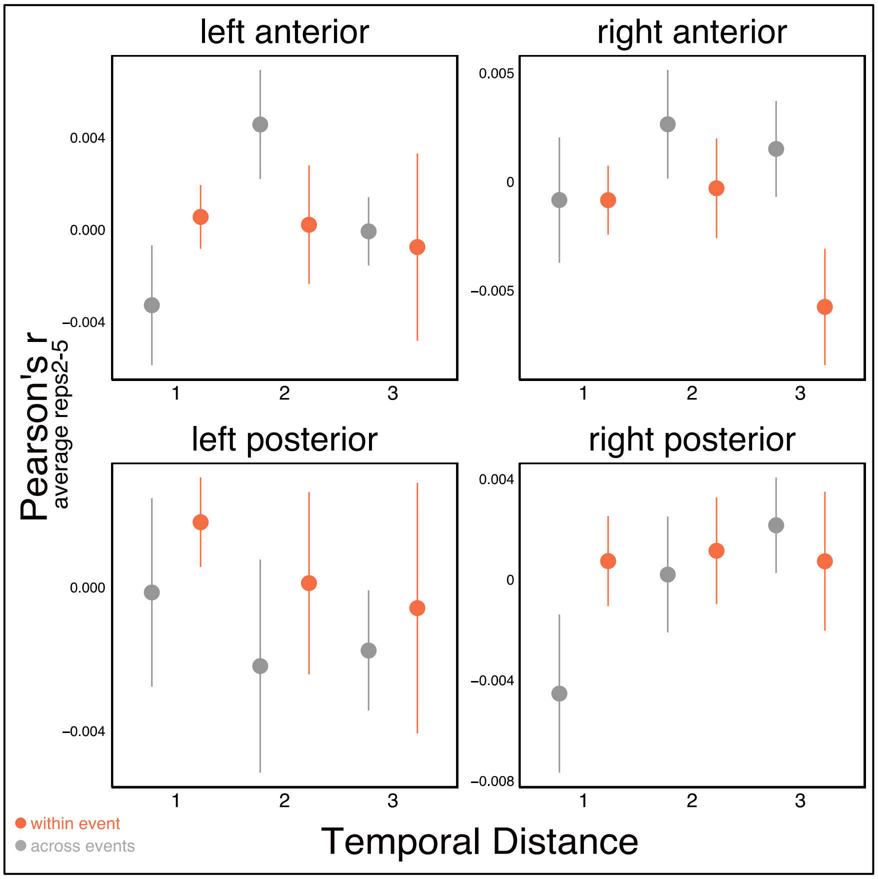


*Supplementary Figure 11.* Similarity values averaged across repetitions 2-5 (after subtracting Repetition1, see Methods), per each ROI (anterior/posterior hippocampus) and in each hemisphere (left/right). Orange: similarity within-event. Grey: similarity across events. Data are presented as mean values, error bars reflect +/- SEM. N = 30. For details and statistical analysis, see text above.

Mixed-effects models formulas

***General notes:***

SubID: participants' numbers

List was treated as factor and was a confound of no interest.

Model comparison was done between the model described as 'Full' and the comparison model ('Comp.').

All models converged without warnings unless specified otherwise. When warning occurred, we reiterated through the model starting from the parameters found in the previous iteration (10 iterations) or trying different optimizers (all optimizers available in the "optimx" package as implemented by the allFit function in dfoptim package in R, with maxfun = 1e5). In our data and models, that did not help. We therefor checked the random effects of the models. When warnings pertained to a boundary singular fit, they stemmed from the random intercepts explaining very little variance (< .0001). Since this is not an issue for our experiment or interpretation, we kept the models as they were. In one case there was a convergence warning (see details below). In all cases, when warning occurred, we further tested the specific hypothesis using a relevant ANOVA or a t-test, and the results were consistent with the mixed-level models, as expected.

***Behavioral data: list learning***

All models included the specification of: family = inverse.gaussian(link = 'identity').

Models testing for interaction of Repetition by Boundary:

Full: scaled-RT ~ Repetition * Boundary + List + (1|subID)

Comp.: scaled-RT ~ Repetition + Boundary + List + (1|subID)

Models testing for the effect of Repetition:

Full: scaled-RT ~ Repetition + Boundary + List + (1|subID)

Comp.: scaled-RT ~ Boundary + List + (1|subID)

Models testing for the effect of Boundary (across all repetitions):

Full: scaled-RT ~ Repetition + Boundary + List + (1|subID)

Comp.: scaled-RT ~ Repetition + List + (1|subID)

Models testing for the effect of Boundary (per each repetition, data of each repetition was included separately):

Full: scaled-RT ~ Boundary + List + (1|subID)

Comp.: scaled-RT ~ List + (1|subID)

For repetition 3 only, the comparison model failed to converge with these warnings:

Model failed to converge with max|grad| = 0.0456116 (tol = 0.002, component 1)

Model is nearly unidentifiable: very large eigenvalue

Reiterating the model and trying different optimizers (as above) did not help. An inspection of the model revealed that this is due to the random intercept explaining little variance (variance < .01). Removing the 'List' confound did not help. Since we were not interested in the random intercepts, and this model was the comparison model, we did not further try to optimize the model and used it as a comparison nevertheless.

In addition, participants' RTs in repetition 3 were averaged based on boundary vs. non-boundary items, and a paired-sample two-tailed t-test was conducted. This revealed a significant boundary effect as well ($t_{(29)} = 5.64$, $p < .0001$).

***Behavioral data: temporal memory test***

All models included the specification of: family = inverse.gaussian(link = 'identity'). The model comparing between Pos1-2 and Pos4-1 did not converge with default settings, so a babyqa optimizer was used by adding 'control = glmerControl(optimizer = "bobyqa")', which solved convergence issues. For consistency, we used this optimizer for all models testing the temporal memory tests. The results did not change if using the default optimizer.

Models testing for the effect of Event Position (as factor), including all positions (Pos1-2/Pos2-3/Pos4-1):

Full: scaled-RT ~ Event Position + List + (1|subID)

Comp.: scaled-RT ~ List + (1|subID)

The models testing the pairwise comparisons (Pos1-2 vs. Pos4-1, Pos2-3 vs. Pos4-1 and Pos1-2 vs. Pos2-3) used the same definitions, but the data only included the data of the relevant event position in each pair of models.

***RSA analyses***

In all models, the effect of Event refers to within vs. across events, and Repetition (1 through 5) was modelled continuously.

Similarity: the similarity values included in the analysis.

The analysis controlling for univariate activation included models identical to those below, with the added term of item1_activation * item2_activation, corresponding to the interaction between the univariate activation of both items contributing to the similarity values (and, automatically, the univariate activation of each of them separately).

***CA3 and DG interaction***

Models testing the interaction of Event by ROI (CA3/DG) by Hemisphere (Left/Right):

Full: Similarity ~ Event * ROI * Hemisphere + Temporal Distance + List + (1|subID)

Comp.: Similarity ~ Event * ROI + ROI * Hemisphere + Event * Hemisphere + Temporal Distance + List + (1|subID)

Models testing the interaction of Event by ROI (CA3/DG) within each hemisphere:

Full: Similarity ~ Event * ROI + Temporal Distance + List + (1|subID)

Comp.: Similarity ~ Event + ROI + Temporal Distance + List + (1|subID)

***Left CA3 analysis***

Models testing the effect of Event:

Full: Similarity ~ Event + Temporal Distance + List + (1|subID)

Comp.: Similarity ~ Temporal Distance + List + (1|subID)

Models testing the effect of Temporal Distance:

Full: Similarity ~ Event + Temporal Distance + List + (1|subID)

Comp.: Similarity ~ Event + List + (1|subID)

Models testing the interaction of Event by Temporal Distance:

Full: Similarity ~ Event * Temporal Distance + List + (1|subID)

Comp.: Similarity ~ Event + Temporal Distance + List + (1|subID)

Models testing the interaction of Repetition by Event by Temporal Distance:

Full: Similarity ~ Repetition * Event * Temporal Distance + List + (1|subID)

Comp.: Similarity ~ Repetition * Event + Repetition * Temporal Distance + Event * Temporal Distance + List + (1|subID)

Models testing the interaction of Repetition by Event:

Full: Similarity ~ Repetition * Event + Temporal Distance + List + (1|subID)

Comp.: Similarity ~ Repetition + Event + Temporal Distance + List + (1|subID)

Models testing the interaction of Repetition by Temporal Distance:

Full: Similarity ~ Repetition * Temporal Distance + Event + List + (1|subID)

Comp.: Similarity ~ Repetition + Temporal Distance + Event + List + (1|subID)

For the models testing the interactions with Repetition, the full and comparison models were on the boundary of a singular fit. For completeness, and although the model did converge, we ran a repeated-measures ANOVA. The data were averaged per participant and per Temporal Distance, Event and Repetition and were entered to a repeated-measures ANOVA with these factors. Consistent with the mixed-level models, a marginally significant interaction of Event by Repetition was observed ($F_{(1,29)} = 3.29$, $p = .08$), whereas other interactions were not significant ($F\text{'s}_{(1,29)} < .2$, $p > .66$).

Models testing the effect of Event in either Repetition2 or Repetition5 (models were the same, used each time with the relevant data):

Full: Similarity ~ Event + Temporal Distance + List + (1|subID)

Comp.: Similarity ~ Temporal Distance + List + (1|subID)

For Repetition 2, both the full and comparison model were on the boundary of a singular fit. A paired-sample two-tailed t-test on the average similarity per participant (within vs. across Event) was not significant ($t_{(29)}$ = .84, $p$ = .41), consistent with the mixed-level model.

Models testing the effect of Color (same/different background color, as a control analysis, only Repetition 5):

Full: Similarity ~ Color + Temporal Distance + List + (1|subID)

Comp.: Similarity ~ Temporal Distance + List + (1|subID)

Both the full and comparison model converged with a boundary singular fit warning. A paired-sample two-tailed t-test on the average similarity per participant (same vs. different Color) was not significant ($t_{(29)}$ = .56, $p$ = .58), consistent with the mixed-level model.

Models testing the effect of CA3 similarity on RT:

Full: Scaled-RT ~ Similarity + Event order + List + (1|subID)

Comp.: Scaled-RT ~ Event order + List + (1|subID)

The same model formulas were used for all analyses described in the section "*CA3 event representations facilitate transitioning between events*", namely, for the main analysis of Neural Clustering predicting Event transition RT, as well as for the follow up analyses.

***Left DG analysis***

Models testing the interaction of Event by Temporal Distance:

Full: Similarity ~ Event * Temporal Distance + List + (1|subID)

Comp.: Similarity ~ Event + Temporal Distance + List + (1|subID)

Models testing the effect of Temporal Distance either within event, or across events (models were the same, used each time with the relevant data):

Full: Similarity ~ Temporal Distance + List + (1|subID)

Comp.: Similarity ~ List + (1|subID)

Models testing the interaction of Repetition by Event by Temporal Distance:

Full: Similarity ~ Repetition * Event * Temporal Distance + List + (1|subID)

Comp.: Similarity ~ Repetition * Event + Repetition * Temporal Distance + Event * Temporal Distance + List + (1|subID)

Models testing interaction of Event by Temporal Distance in either Repetition2 or Repetition5 (models were the same, used each time with the relevant data):

Full: Similarity ~ Event + Temporal Distance + List + (1|subID)

Comp.: Similarity ~ Temporal Distance + List + (1|subID)

These models converged with a boundary singular fit warning. A repeated-measures ANOVA with data averaged per participant (and per Event and Temporal Distance conditions) yielded consistent results with the mixed-level models. In Repetition 5, a significant Event by Temporal Distance was observed ($F_{(1,29)} = 7.08$, $p = .013$), but not in Repetition 2 ($F_{(1,29)} = .06$, $p = .81$).

Models testing the effect of Temporal Distance within event or across events in Repetition 5, testing all lags, or lags of 1 vs. 2 or 1 vs. 3 (models were the same, used each time with the relevant data):

Full: Similarity ~ Temporal Distance + List + (1|subID)

Comp.: Similarity ~ List + (1|subID)

The models testing all lags, as well as lag of 1 vs. 2, converged with a boundary singular fit warning. Data were averaged per participant and per Temporal Distance condition (within- or across events separately). Consistent with the mixed-level model, a one-way repeated measures ANOVA testing for the main effect of Temporal Distance yielded a main effect within event ($F_{(1,29)}$ = 5.53, $p$ = . 026), but not across events ($F_{(1,29)}$ = .08, $p$ = . 78). Likewise, paired-sample two-tailed t-tests yielded a significant effect of Temporal Distance comparing lag of 1 vs. 2 within event ($t_{(29)}$ = 2.38, $p$ = .02), but not across events ($t_{(29)}$ = .50 $p$ = .62).

Models testing the effect of Event within each lag in Repetition5 (models were the same, used each time with the relevant data):

Full: Similarity ~ Event + List + (1|subID)

Comp.: Similarity ~ List + (1|subID)

The models testing lag 1 and 2 converged with a boundary singular fit warning. Consistent with the mixed-level model, paired-sample two-tailed t-tests on data averaged per participant and per within vs. across Event (in each lag separately), resulted in a significant effect of Event for lag 1 ($t_{(29)}$ = 2.49, $p$ = .02), but not lag 2 ($t_{(29)}$ = 1.066, $p$ = .29).

*References*

Brunec, I. K., Bellana, B., Ozubko, J. D., Man, V., Robin, J., Liu, Z.-X., Grady, C., Rosenbaum, R. S., Winocur, G., Barense, M. D., & Moscovitch, M. (2018). Multiple Scales of Representation along the Hippocampal Anteroposterior Axis in Humans. *Curr. Biol.*, *28*(13), 2129-2135.e6. https://doi.org/10.1016/j.cub.2018.05.016

Chawla, M. K., Guzowski, J. F., Ramirez-Amaya, V., Lipa, P., Hoffman, K. L., Marriott, L. K., Worley, P. F., McNaughton, B. L., & Barnes, C. A. (2005). Sparse, environmentally selective expression of Arc RNA in the upper blade of the rodent fascia dentata by brief spatial experience. *Hippocampus*, *15*(5), 579–586. https://doi.org/10.1002/hipo.20091

Danker, J. F., & Anderson, J. R. (2010). The ghosts of brain states past: Remembering reactivates the brain regions engaged during encoding. *Psychol. Bull.*, *136*(1), 87–102. https://doi.org/10.1037/a0017937

Grill-Spector, K., Henson, R., & Martin, A. (2006). Repetition and the brain: Neural models of stimulus-specific effects. *Trends in Cognitive Sciences*, *10*(1), 14–23. https://doi.org/10.1016/j.tics.2005.11.006

Jung, M. W., & McNaughton, B. L. (1993). Spatial selectivity of unit activity in the hippocampal granular layer. *Hippocampus*, *3*(2), 165–182. https://doi.org/10.1002/hipo.450030209

Madar, A. D., Ewell, L. A., & Jones, M. V. (2019). Temporal pattern separation in hippocampal neurons through multiplexed neural codes. *PLoS Comput. Biol.*, *15*(4), e1006932. https://doi.org/10.1371/journal.pcbi.1006932

Poppenk, J., & Moscovitch, M. (2011). A hippocampal marker of recollection memory ability among healthy young adults: Contributions of posterior and anterior segments. *Neuron*, *72*(6), 931–937. https://doi.org/10.1016/j.neuron.2011.10.014

Ritchey, M., Wing, E. A., LaBar, K. S., & Cabeza, R. (2013). Neural similarity between encoding and retrieval is related to memory via hippocampal interactions. *Cereb. Cortex*, *23*(12), 2818–2828. https://doi.org/10.1093/cercor/bhs258

Thorp, J. N., Gasser, C., Blessing, E., & Davachi, L. (2022). Data-Driven Clustering of Functional Signals Reveals Gradients in Processing Both within the Anterior Hippocampus and across Its Long Axis. *The Journal of Neuroscience*, *42*(39), 7431–7441. https://doi.org/10.1523/JNEUROSCI.0269-22.2022

Tompary, A., Duncan, K., & Davachi, L. (2016). High-resolution investigation of memory-specific reinstatement in the hippocampus and perirhinal cortex. *Hippocampus*, *26*(8), 995–1007. https://doi.org/10.1002/hipo.22582

Tompary, A., & Davachi, L. (2017). Consolidation Promotes the Emergence of Representational Overlap in the Hippocampus and Medial Prefrontal Cortex. *Neuron*, *96*(1), 228–241. https://doi.org/10.1016/j.neuron.2017.09.005
